# MitoQuicLy: a high-throughput method for quantifying cell-free DNA from human plasma, serum, and saliva

**DOI:** 10.1101/2023.01.04.522744

**Authors:** Jeremy Michelson, Shannon Rausser, Amanda Peng, Temmie Yu, Gabriel Sturm, Caroline Trumpff, Brett A. Kaufman, Alex J. Rai, Martin Picard

**Affiliations:** Department of Psychiatry, Division of Behavioral Medicine, Columbia University Irving Medical Center, New York, USA; Department of Biochemistry and Biophysics, University of California, San Francisco, San Francisco, California, USA; Center for Metabolism and Mitochondrial Medicine and the Vascular Medicine Institute, Division of Cardiology, Department of Medicine, University of Pittsburgh School of Medicine; Department of Pathology and Cell Biology, Columbia University Irving Medical Center, New York, NY, USA; Department of Neurology, H. Houston Merritt Center, Columbia Translational Neuroscience Initiative, Columbia University Irving Medical Center, New York, NY, USA; New York State Psychiatric Institute, New York, NY, USA

**Author notes:** **Corresponding author:** 1051 Riverside Drive, Kolb 4, New York, NY 10032, United States.

**Keywords:** mitochondria, circulating nucleic acids, DNA isolation, mitochondrial DNA, protocol, cell-free DNA

## Abstract

Circulating cell-free mitochondrial DNA (cf-mtDNA) is an emerging biomarker of psychobiological stress and disease which predicts mortality and is associated with various disease states. To evaluate the contribution of cf-mtDNA to health and disease states, standardized high-throughput procedures are needed to quantify cf-mtDNA in relevant biofluids. Here, we describe MitoQuicLy: <u>Mito</u>chondrial DNA <u>Qu</u>antification in <u>c</u>ell-free samples by <u>Ly</u>sis. We demonstrate high agreement between MitoQuicLy and the commonly used column-based method, although MitoQuicLy is faster, cheaper, and requires a smaller input sample volume. Using 10 µL of input volume with MitoQuicLy, we quantify cf-mtDNA levels from three commonly used plasma tube types, two serum tube types, and saliva. We detect, as expected, significant inter-individual differences in cf-mtDNA across different biofluids. However, cf-mtDNA levels between concurrently collected plasma, serum, and saliva from the same individual differ on average by up to two orders of magnitude and are poorly correlated with one another, pointing to different cf-mtDNA biology or regulation between commonly used biofluids in clinical and research settings. Moreover, in a small sample of healthy women and men (n=34), we show that blood and saliva cf-mtDNAs correlate with clinical biomarkers differently depending on the sample used. The biological divergences revealed between biofluids, together with the lysis-based, cost-effective, and scalable MitoQuicLy protocol for biofluid cf-mtDNA quantification, provide a foundation to examine the biological origin and significance of cf-mtDNA to human health.

## 1. Introduction

Circulating cell-free nucleic acids (cf-DNA and cf-RNA) have emerged as important biomarkers in medicine and in research (Pös et al., 2018). In the clinic, the concentration and sequence of circulating DNA are used as a prognostic and diagnostic biomarker in oncology and non-invasive prenatal testing (Norton et al., 2015; Otandault et al., 2019). Circulating DNA levels are also persistently elevated in various pathologies, including sepsis, cancer, and trauma (Dwivedi et al., 2012; Faust et al., 2020; Nakahira et al., 2013; Schwarzenbach et al., 2011). A substantial fraction of circulating DNA originates from mitochondria, released as circulating cell-free mitochondrial DNA (cf-mtDNA). cf-mtDNA concentration (copies per µl) is selectively elevated in certain disease states, and plasma cf-mtDNA is prospectively associated with a 4-8 fold increased mortality risk among critically ill patients (Nakahira et al., 2013) and predicts mortality among sepsis patients (Kung et al., 2012). Some studies, although not all (Daniels et al., 2022), have also found associations between cf-mtDNA levels and inflammatory processes (Pinti et al., 2014; Zhang et al., 2010) (for a comprehensive review, see (Boyapati et al., 2017)). However, in many contexts, cf-mtDNA likely has non-inflammatory roles possibly related to the inter-cellular transfer of organelles and nucleic acids (Brestoff et al., 2021; Levoux et al., 2021; Miliotis et al., 2019; Trumpff et al., 2021). Together these findings position cf-mtDNA as an emerging yet poorly understood disease biomarker.

While the cellular origins and physiological functions of cf-mtDNA are not yet clear, cf-mtDNA is likely selectively released. In the plasma of healthy individuals, there are approximately 50,000 times more copies of cf-mtDNA than cf-nDNA (Meddeb et al., 2019a). However, there are only about ∼200-5,000 copies of mtDNA per cell in most tissues (Picard, 2021; Wachsmuth et al., 2016), which would be the default ratio if genomic material was indiscriminately released from dying cells. Moreover, certain insults, including viral infection, sepsis, and psychological stress elevate blood cf-mtDNA – but not cf-nDNA (e.g., (Trumpff et al., 2019)), suggesting that selective release of mtDNA occurs without equivalent nDNA release.

Regarding the form of transport of cf-mtDNA in biofluids, in healthy human plasma, most cf-mtDNA pellets with centrifugation and can be removed by a 0.22 µm filter, and therefore appears to be membrane-enclosed, likely contained within intact mitochondria or potentially mitochondria derived vesicles (Al Amir Dache et al., 2020; D’Acunzo et al., 2021; Song et al., 2020; Stephens et al., 2020). Thus, both the selective release of the mitochondrial genome and the pathophysiological relevance of cf-mtDNA outlined above call for studies examining the significance of cf-mtDNA as a biomarker.

Like many biomarkers that are responsive to environmental and physiological exposures, and which therefore must be examined at high frequency or in large sample sizes to decipher the signal from noise, cf-mtDNA needs to be examined across biofluids, time scales, and in diverse populations, making the throughput of cf-mtDNA assays an important assay parameter. For instance, plasma cf-mtDNA levels may increase over decades with human aging (Pinti et al., 2014), blood cf-mtDNA levels changes over weeks to months during pregnancy (serum) (Cushen et al., 2020), with exercise training (serum) (Nasi et al., 2016), and following psychotropic treatment (plasma) (Lindqvist et al., 2018). Moreover, over much shorter time scales, studies in plasma (Hummel et al., 2018; Lindqvist et al., 2016; Shockett et al., 2016; Stawski et al., 2017), serum (Trumpff et al., 2019), and saliva (Trumpff et al., 2022) show that cf-mtDNA concentrations change several-fold over periods ranging from minutes to hours (reviewed in (Trumpff et al., 2021)), calling for longitudinal and prospective studies with repeated measures to accurately capture cf-mtDNA dynamics. Thus, the need to examine cf-mtDNA across varying timescales and broadly representative participant populations (Clayton and Collins, 2014) calls for studies with large sample sizes, compounding the need for high-throughput, cost-effective, scalable laboratory methods.

Remarkable progress has been made in developing protocols to collect, isolate, and quantify blood cf-DNA (Cox et al., 2020; El Messaoudi et al., 2013; Meddeb et al., 2019a; Meddeb et al., 2019b; Trumpff et al., 2021; Vishwakarma et al., 2022; Ware et al., 2020). However, methods used across studies still vary substantially between laboratories and across biofluid types, including different types of anticoagulated plasma types, serum types, saliva, and others (Trumpff et al., 2021). The adoption of standard methodologies offering portability and scalability across biofluids and sample sizes rests on the availability of scalable, cost-effective methods that require standard laboratory equipment. To that end, we developed *MitoQuicLy*: <u>Mito</u>chondrial DNA <u>qu</u>antification in <u>c</u>ell-free samples via <u>ly</u>sis. MitoQuicLy is a high-throughput, lysis-based DNA quantification method similar to what has been described previously to quantify mtDNA copy number in single cells (Grünewald et al., 2016), cell mixtures (Longchamps et al., 2020), and tissue samples (Picard et al., 2012). MitoQuicLy uses commonly available laboratory equipment and consumables, small sample input volume, and inexpensive reagents with minimal hands-on time. Here we describe the development and validation of MitoQuicLy and deploy this method in commonly used blood and saliva biofluid types to provide foundational knowledge for biomarker studies, and to inform future mechanistic studies required to determine the nature and physiological functions of cf-mtDNA in humans.

## 2. Materials and Methods

MitoQuicLy was developed and tested using plasma, serum, and saliva from healthy participants recruited in two cohorts. In the first cohort, only EDTA plasma was collected. In the second cohort, we collected plasma in EDTA, heparin, and citrate tubes; serum in red top (clot activator) and gold top tubes (clot activator + serum separator gel); and saliva using salivettes (see below for details). All participants provided written informed consent for their participation and publication of the results. This study was approved by New York State Psychiatric Institute Institutional Review Board protocol #7618.

### 2.1. Cohort 1: blood collection and cell-free plasma preparation

Blood was collected from 10 healthy participants by venipuncture with a 21-gauge needle (BD #367281). The first 9 mL of blood drawn was used for other biomarker measurements. Blood was then drawn into 10 mL K_2_EDTA blood collection tubes (BD #366643). The tube was inverted 10-12 times to ensure thorough mixing. To immediately separate plasma from the major cellular constituents that may contain and/or release mtDNA (Aucamp et al., 2018; Boudreau et al., 2014), blood was immediately centrifuged at 1,000g for 5 minutes at room temperature with no brake. Samples were then placed on ice and remained at 4ºC for the remainder of the sample processing. After 30 minutes, the K_2_EDTA tubes were centrifuged again at 2,000g for 10 minutes at 4ºC. Approximately 80% of the plasma was drawn from the top of the tube and placed in a new 15 mL conical tube. To deplete platelets and other cellular debris, plasma was again centrifuged at 2,000g for 10 minutes at 4ºC. 90% of the plasma supernatant was transferred to a new 15 mL conical tube and mixed by inversion. The cell-free plasma was divided into 0.5 mL aliquots into 1.5 mL Eppendorf tubes and stored at -80ºC.

### 2.2. Cohort 2: blood collection and cell-free sample preparation

To validate MitoQuicLy and compare it across various sample types, we recruited 34 healthy adults from the university and surrounding community. Participants were recruited via flyers posted on message boards and circulated via email lists. Informed consent was obtained via video conference or telephone call, and informed consent was obtained remotely and stored electronically in the Research Electronic Data Capture (REDCap) database. Demographic information was collected through an electronic questionnaire.

Before the blood draw, participants sat in a chair and rested while equipped with an electronic blood pressure monitor. Measures of blood pressure and heart rate were obtained from the average of two measurements. Venipuncture was performed with a 21-gauge needle (BD #367281), and 66 mL of blood was collected into 13 tubes. Blood was drawn between 08:30 and 14:30, and participants were not required to fast. Blood samples were drawn consecutively in the following order: Citrate (2 × 2.7 mL), Heparin (6.0 mL), EDTA (6.0 mL), Red top serum (6.0 mL), and Gold top serum tubes (6.0mL). After collection, all tubes were inverted 10-12 times to ensure thorough mixing. Concurrently, saliva was sampled using a cotton swab Salivette (Sarstedt #51.1534.500) on the tongue for 5 minutes, beginning when blood started to flow into collection tubes.

Plasma was prepared from 2 × 2.7 mL blue top citrate tubes, 1 × 6.0 mL green top heparin tube, and 1 × 6.0 mL lavender top EDTA tube. Serum was prepared from a 6.0 mL red top serum tube and a 6.0 mL gold top serum separation tube. Plasma samples were immediately centrifuged at 1,000g for 5 minutes at room temperature. Serum samples were incubated at room temperature for 30 minutes before centrifugation at 1,000g for 5 minutes (red top serum) or 1,000g for 10 minutes (gold top serum). After this initial step, all samples were treated identically. To avoid cellular contamination, the plasma/serum supernatant was drawn from the top of the tube, leaving about 1 cm excess remaining above the buffy coat. To pellet and remove the majority of remaining platelets, plasma/serum was transferred into 1.5 mL Eppendorf tubes and centrifuged again at 5,000g for 10 minutes at room temperature. The clean plasma/serum supernatant was transferred to a new tube, leaving about 100 µL above the platelet pellet. The resulting clean plasma/serum was mixed by pipetting and aliquoted in 150 µL aliquots. Aliquots were frozen at -80ºC.

Salivettes were immediately centrifuged at 1,000g for 5 minutes at room temperature. To avoid cellular contamination with leukocytes or epithelial buccal cells (Theda et al., 2018), the saliva supernatant was drawn from the top of the tube, leaving about 200 µl above the cell pellet. Saliva was transferred to 1.5 mL Eppendorf tubes and stored at -80°C. Before use, saliva was thawed and centrifuged at 5,000g for 10 minutes, and supernatant cf-DNA was quantified using MitoQuicLy. All assays were performed within one year of sample collection.

### 2.3. MitoQuicLy method

A schematic of the MitoQuicLy protocol is shown in Figure 1A. Once cell- and platelet-depleted biofluids (plasma, serum, or saliva) are obtained, 10 µL of sample is added to 190 µL lysis buffer in a standard 96-well semi-skirted PCR plate. The lysis buffer is composed of 6% Tween-20 (Sigma #P1379), 114 mM Tris-HCl pH 8.5 (Sigma #T3253), and 200 µg/mL Proteinase K (Thermofisher #AM2548). The plate wells are sealed with PCR strip caps, and the samples are mixed by vortexing followed by a brief pulse spin. The sample/lysis buffer mixture is then incubated overnight at 55ºC for 16h in a standard thermocycler, followed by Proteinase K inactivation at 95ºC for 10 minutes, and held at 4ºC until qPCR (Supplemental Figure 1E). The lysate can then be used directly in qPCR assays or stored at -80ºC for later use. Figure 1B illustrates the lysis reaction, highlighting the complimentary proteolytic and lipid-dissolving actions of the protease and detergent, respectively – yielding accessible DNA for quantification. The detailed, step-by-step MitoQuicLy protocol is available in Supplemental File 1.

**Figure 1:**
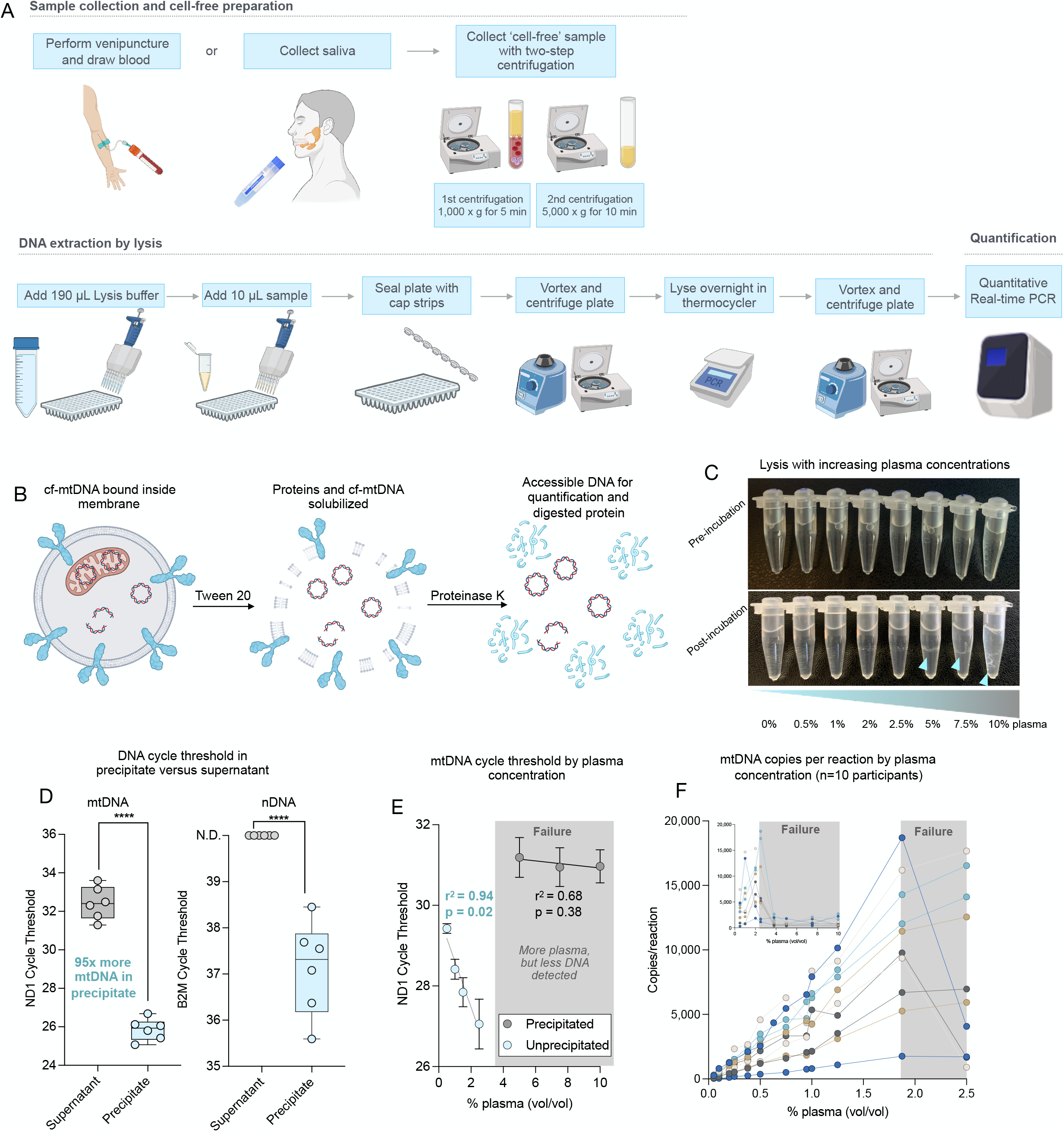
Optimizing collection, lysis and qPCR protocol. (**A**) Schematic of MitoQuicLy Protocol. Biological samples are collected (min 5 mL) and centrifuged twice to remove cells. Cell-free samples are added to lysis buffer in a 96-well plate, sealed with cap strips and incubated overnight in a standard thermocycler before quantification using qPCR. (**B**) Schematic of lysis reaction. Tween-20 solubilizes membranes and Proteinase K digests proteins. The lysate is used directly as template DNA for qPCR. (**C**) Pooled EDTA plasma was lysed at a range of concentrations. At higher concentrations (≥5%), a stringy white precipitate forms (blue arrows). (**D**) DNA was quantified from either the supernatant (grey) or the precipitate (blue), indicating the high abundance of DNA in the precipitate. No nDNA was detected in supernatants (Ct undetermined: >40), but nDNA was detected in precipitates albeit at a high Ct. N.D.: Not detected. **** p<0.0001, unpaired t test. (**E**) Samples with precipitation have less DNA than unprecipitated samples. Plasma was lysed in a dilution series from 10% to 0.5%. A linear relationship between cycle threshold and plasma concentration was observed below 2.5%, but not above 2.5%; simple linear regression. (**F**) Dilution series from 10 participants lysed at a series of concentrations from 10% to 0.5% (inset) and at concentrations from 2.5% to 0.05%, confirming the assay breakpoint, and linearity below 2%. Each participant datapoint was measured in triplicates.

For validation purposes, we used Qiagen DNeasy Blood and Tissue Kit (Qiagen #69504) as our Silica Membrane Kit (column) comparison following the manufacturer’s instructions for non-nucleated blood. We combined 100 µL of plasma or serum, 20 µL Proteinase K, 100 µL PBS and 200 µL of Buffer AL. Samples were thoroughly vortexed before being incubated at 56ºC for 10 minutes. 200 µL pure ethanol was added, and the sample was mixed by vortex. The resulting lysate was pipetted onto a DNeasy Mini spin column and centrifuged at 6,000g for 1 minute. The column was washed with 500 µL Buffer AW1, then 500 µL Buffer AW2, and finally, DNA was eluted in 200 µL of buffer AE.

### 2.4. DNA standards

Primary human dermal fibroblasts (hFB1) were obtained with informed consent from a healthy 29-year-old male donor (IRB #AAAB0483). Fibroblasts were grown to confluence in physiological glucose DMEM supplemented with 10% FBS, 1% non-essential amino acids, 50 µg/mL uridine, and 10 µM Palmitate in a T175 flask as described in (Sturm et al., 2022). Cells were trypsinized and counted using a Countess II Automated Cell Counter (Thermofisher #A27977), yielding two pellets of 12-14 million cells. Each pellet was lysed in 2 mL of lysis buffer (described above) following the same procedure as for biofluids. Total DNA was quantified by Qubit. From each pellet, 8.38-8.58 µg of DNA was extracted (yield: 652 ng of DNA per million cells). The two lysates were combined and diluted to a stock concentration of 1 µg/mL, aliquoted and frozen at -80ºC until use.

### 2.5. Digital droplet PCR

The absolute number of mtDNA and nDNA copies in the hFB1 DNA standard was quantified by digital PCR (dPCR). The DNA was serially diluted 1:4 with nuclease-free water. Standard dilutions were combined with QuantStudio Digital PCR Mastermix, TaqMan primers and probes for ND1 (mtDNA) and B2M (single-copy nDNA gene). Primer and probe sequences are available in Supplemental File 1. The final concentrations of primers were 300 nM and the final concentrations of probes were 100 nM. The dPCR reaction mixture was loaded onto QuantStudio 3D Digital PCR 20K Chip V2 (Thermofisher) as per the manufacturer’s instructions. PCR was performed on a standard thermocycler (ProFlex, Thermofisher) with the following conditions: 96ºC for 10 min, followed by 39 cycles of 60ºC for 2 min and 98ºC for 30 seconds, 60ºC for 2 min, and held at 10ºC until the chip was read. For ND1, the sixth dilution (i.e., 0.97 ng/mL) contained 6259.1 copies/reaction. For B2M, the second dilution (i.e., 625 ng/mL) contained 1470.2 copies/reaction. From these results, we calculated the number of copies per reaction for other dilutions, computed the natural logarithm of each dilution’s copies/reaction, and thereafter included 8 serial dilutions of the hFB1 standards at 1:4 on each qPCR plate. The correlation coefficient between the natural logarithm of each dilution’s copies/reaction with the cycle thresholds was r^2^=0.99. The slope and intercepts of the standard curves were then used to compute copies/reaction for each experimental biofluid sample.

### 2.6. Quantitative real-time PCR

cf-mtDNA and cf-nDNA concentration was quantified by duplex quantitative polymerase chain reaction for ND1 (mtDNA) and B2M (nDNA) (see Supplemental File 1 for details of the primers/probe sequences). The ND1 amplicon is 69 bp and targets the mtND1 gene (ENSG00000198888.2) in the mitochondrial genome from position 3,485-3,553. The B2M amplicon is 96 bp and targets an exon in the B2M gene (ENSG00000166710.22) in chromosome 15 from position 44,715,455-44,715,550. *In silico* specificity was assessed using the UCSC genome browser, which confirmed that these primers only target one genomic location. Primers and probes were synthesized by idtDNA and purified by standard techniques (standard desalting for primers, HPLC for probes). For each experimental plate, a mastermix was prepared by combining 2xTaqMan Universal Mastermix Fast (Life Technologies #4444965) with 10 µM ND1 Primers F+R, 10 µM B2M Primers F+R, 5 µM ND1 Probes-VIC, and 5 µM B2M Probes-FAM (idtDNA.com). The final concentrations of primers were 300nM and the final concentrations of probes were 100nM.

An automated repeater pipette dispensed 12 µL of mastermix into each well of the 384-well qPCR reaction plate (ThermoFisher # 4309849). Next, 8 µL of lysate (input DNA) from the lysis plate was added using a multichannel pipette for a final reaction volume of 20 µL. All reactions were carried out in triplicates. The plates were sealed with MicroAmp optical adhesive film (ThermoFisher #4311971) and briefly pulse-spun to 1,000g. qPCR was performed on the QuantStudio 7 Flex Real-Time PCR System (ThermoFisher #4485701) with the following cycling conditions: 50ºC for 2 minutes, 95ºC for 20 seconds, followed by 40 cycles of 95ºC for 1 second, and 60ºC for 20 minutes (total runtime: 40 minutes) (Supplemental Figure 1D). A Δ normalized reaction (ΔRn) threshold of 0.08 was used to determine cycle thresholds for both ND1 and B2M using QuantStudio Real-Time qPCR software. Averages, standard deviations, and coefficients of variation (CV) were computed between qPCR triplicates. Outlier wells that resulted in CVs greater than 2% between triplicates were discarded.

### 2.7. qPCR validation

To establish the detection limit in our qPCR assay, 16 serial 1:4 dilutions of hFB1 DNA standards in nuclease-free water were performed (first concentration = 0.25 µg/mL) and were subjected to qPCR. ND1 was detectable in dilutions 1-10, while B2M was detectable only for dilutions 1-6, consistent with the greater copies of mtDNA relative to nDNA for this cell line (Sturm et al., 2021) (Supplemental Figure 1A and 1B). Plotting the log_10_ of copy number against the cycle threshold for each standard dilution yielded r^2^ values >0.99 for ND1 and B2M, and qPCR efficiency (Supplemental Figure 1C) was 102.48% for ND1 (slope=-3.264) and 90.53% for B2M (slope=-3.572). According to the computed copies/reaction for each dilution obtained from digital PCR, the detection limit for ND1 was 6.1 copies/reaction, while the detection limit for B2M was 1.4 copies/reaction. The average Ct for a No Template Control was 35.80 for ND1 and undetermined for B2M.

### 2.8. Statistical Analyses

Statistical analyses were performed using Prism 9 statistical analysis software. All data, analyses, and graphs are included in Supplemental File 2. Supernatant and precipitate cycle thresholds were compared with unpaired t tests (Figure 1D). The relationship between sample percent (vol/vol) and cycle threshold was evaluated using simple linear regression (Figure 1E). When we modified buffer component concentrations, optimal concentrations were chosen by calculating the modal maximum for each participant (Supplemental Figure 2). Two-component buffers were evaluated using a 2-way ANOVA comparing each modified buffer with the LLL buffer using the Dunnett correction for multiple comparisons (Figure 2). Spearman’s correlations were computed for the correlation of sample percent and cf-DNA measured using the HHL buffer (Figure 3). When calculating the variability of 96 replicates of MitoQuicLy, histograms were created with bins sized to 1/15^th^ the range between maximum and minimum values.

**Figure 2:**
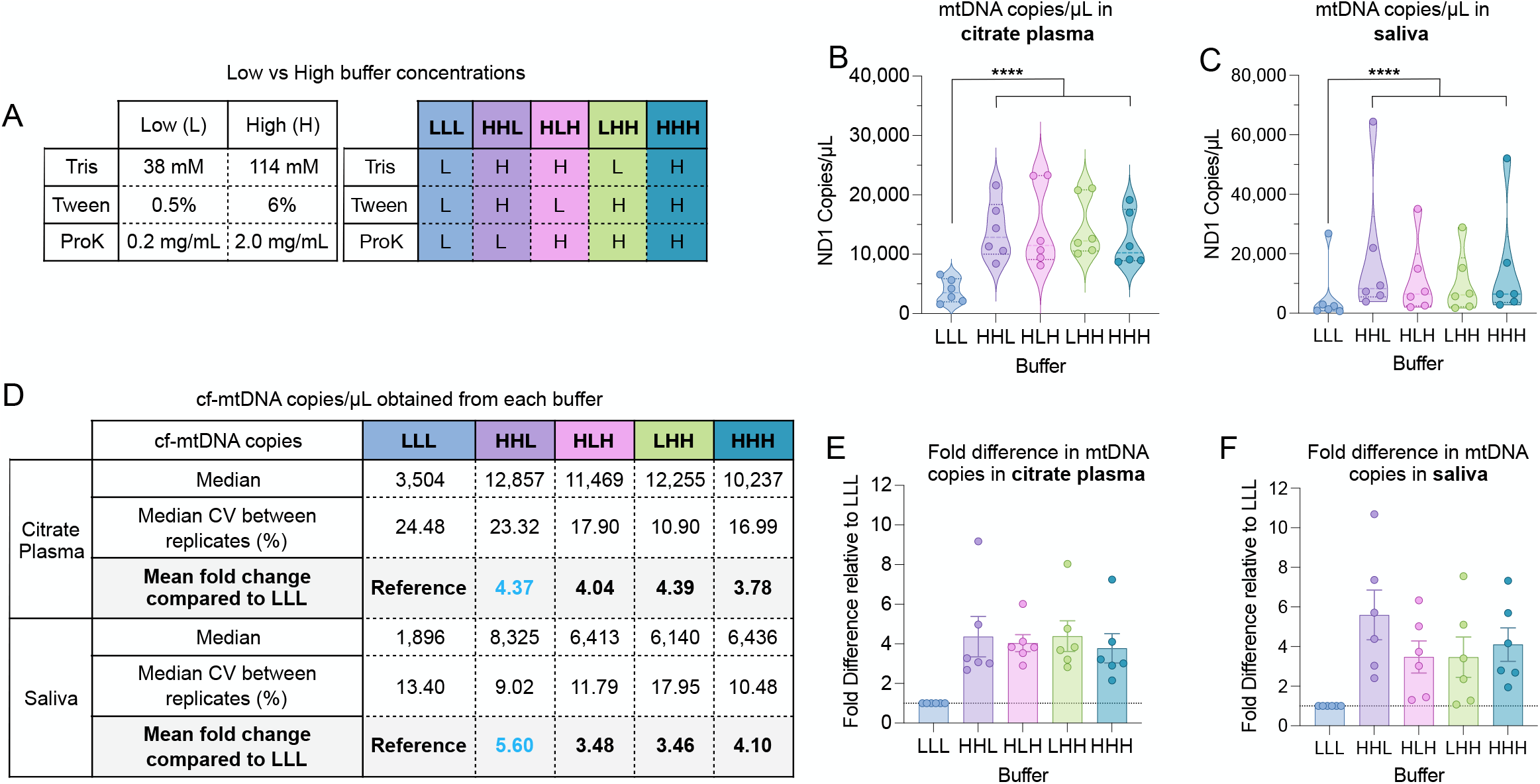
Modifying buffer component concentrations. (**A**) A range of buffer compositions were tested (see supplement) to compare low (L) and high (H) buffer component, yielding 5 different buffer combinations tested against the original LLL (using the modal maximum values in single buffer experiments). (**B and C**) Citrate plasma and saliva cf-mtDNA levels measured in matching sample types from n=6 individuals, indicating improved yield with revised buffer combinations. Each datapoint was measured in triplicates. (**D**) mtDNA copies detected for each buffer combination relative to LLL. (**E and F**) Fold-change in cf-mtDNA yield relative to the LLL buffer, revealing HHL as optimal for citrate plasma and saliva. n=6 individuals, 3 women, 3 men; p<0.05*, p<0.01**, p<0.001***, p<0.0001****, two-way ANOVA.

**Figure 3:**
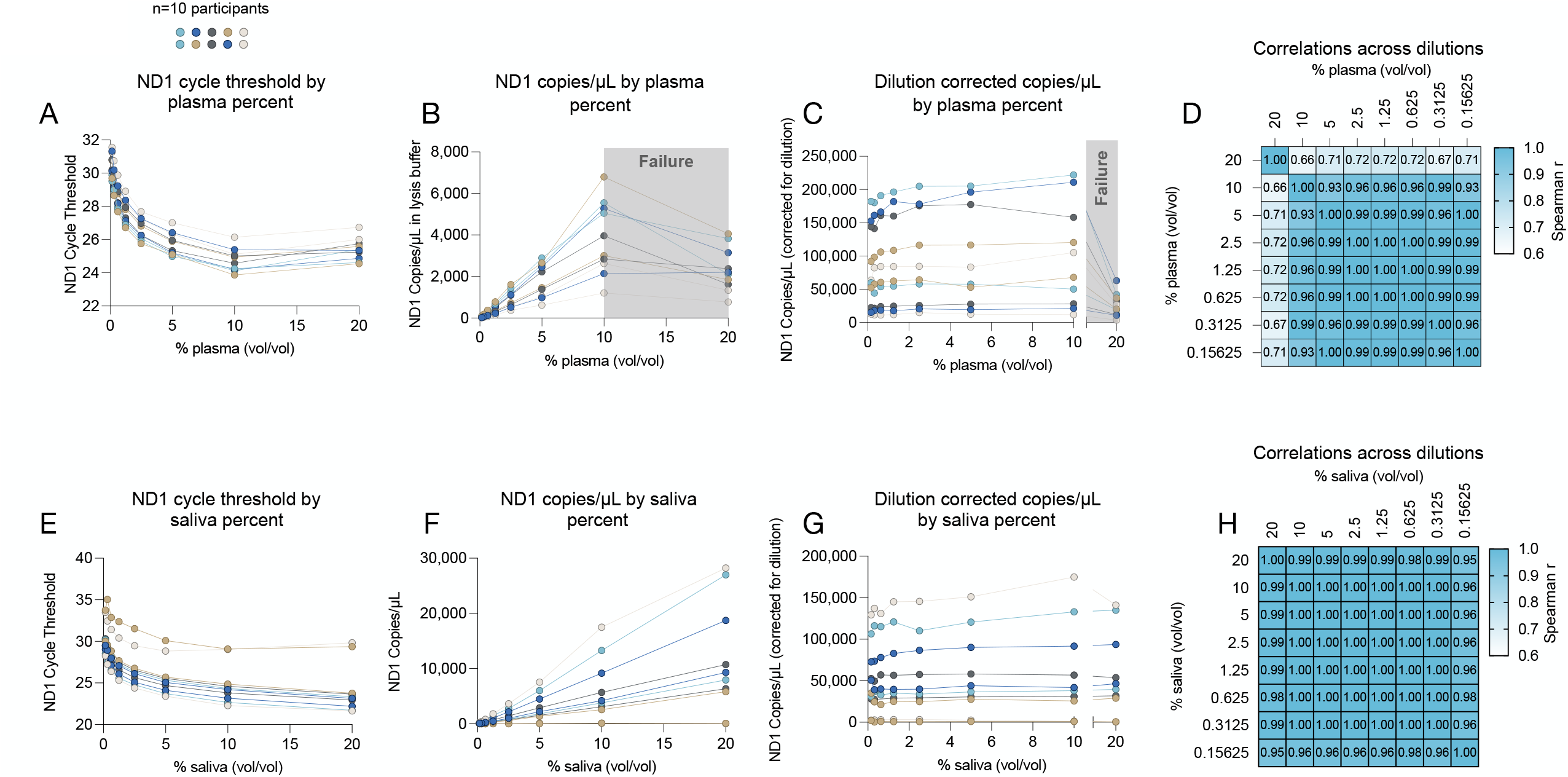
Titration of sample concentration with optimized MitoQuicLy buffer. (**A and E**) ND1 quantification in citrate plasma and saliva from n=10 people lysed in eight 2-fold serial dilution ranging from 20% to 0.078% (vol/vol). Each person’s serial dilution is represented by points on a connected line. (**B and F**) Copies/µL derived from a standard curve, showing that the assay in citrate plasma is linear from 0 to 10% plasma, but “breaks” between 10 and 20% (grey shading). In saliva, the assay range is linear across all concentrations tested. (**C and G**) Dilution-corrected copies/µL in plasma and saliva. From 0-10% plasma or saliva, all samples are reliably rank-ordered at any of the titrated concentrations. (**D and H**) Spearman’s r for plasma and saliva dilution curves (copies/µL) illustrating high correlations between all dilution ≤10%. n=10 individuals, 5 women, 5 men.

Descriptive statistics were computed to obtain CV between lysis replicates (Figure 4). To compare MitoQuicLy with column extraction, we used simple linear regression for the whole dataset (Figure 5A) and each sample type (Figure 5B). Differences in ND1 Cts between sample types were evaluated using a mixed-effects model with multiple comparisons using Bonferroni correction, as we were missing saliva samples from 4/34 people (Figure 6B). Correlations between each biofluid ND1 Ct were evaluated using non-parametric Spearman’s rank correlation (Figure 6D). For exploratory analyses of sex differences, we used Hedges’ g as a measure of effect size (mean ND1 copies relative to the pooled and weighted standard deviation, Figure 6E), which, unlike Cohen’s d, includes a correction for small sample size (Goulet-Pelletier and Cousineau, 2018). A Hedge’s g value of ±1.0 indicates that sexes differ by one standard deviation. Positive values indicate levels higher in women; negative values indicate levels higher in men. To evaluate the relationship between cf-mtDNA levels and blood biomarkers, we used non-parametric Spearman’s correlation with two-tailed alpha set at 0.05 (Figure 7). Statistical significance was set at p<0.05.

**Figure 4:**
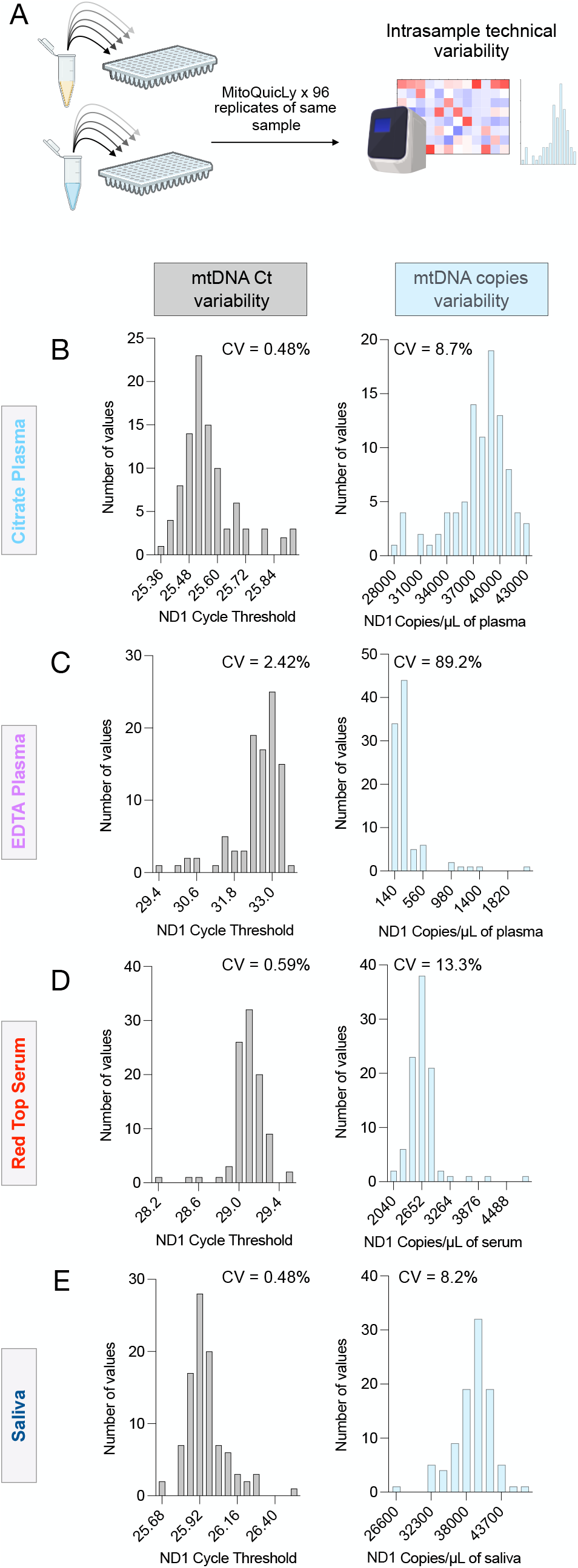
MitoQuicLy variability. (**A**) cf-mtDNA quantified on pooled samples from 34 healthy controls (14 women, 20 men) were run as 96 technical replicates using MitoQuicLy. Frequency distributions of cycle thresholds (Ct, *left*) or copies/µL (*right*) for citrate plasma (**B**), EDTA plasma (**C**), serum (**D**), and saliva (**G**). CVs were acceptable for all samples except EDTA plasma, likely due to the low DNA concentration in EDTA plasma.

**Figure 5:**
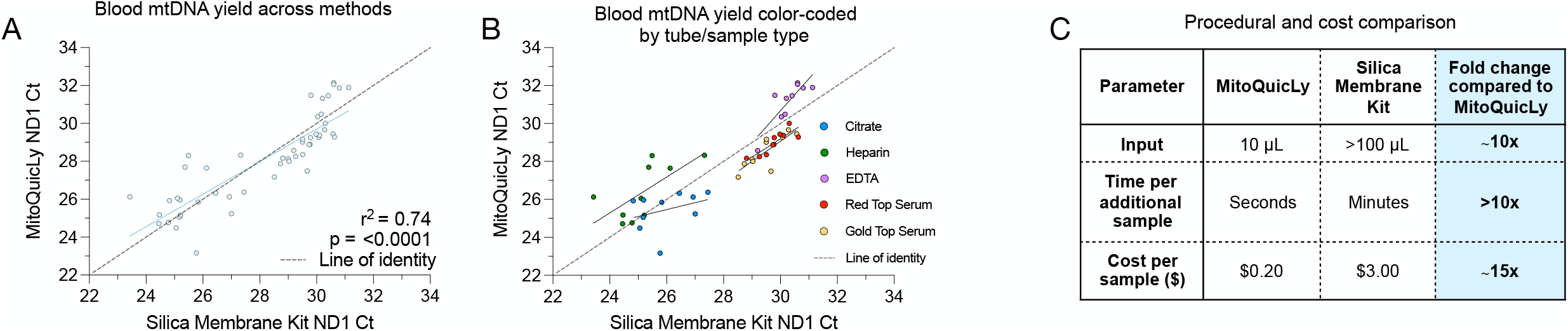
Comparison of cf-mtDNA levels between MitoQuicLy and silica-based membrane kit. (**A**) cf-DNA quantified from various plasma or serum types using either MitoQuicLy or a silica-based membrane column DNA extraction kits (DNeasy, Qiagen). A total of 50 samples were analyzed (5 biofluids, n=10 individuals [5 women, 5 men]). (**B**) Data from (A) color-coded by sample type. (**C**) Procedural and financial comparison of key parameters between MitoQuicLy and standard silica-based membrane DNA extraction kits.

**Figure 6:**
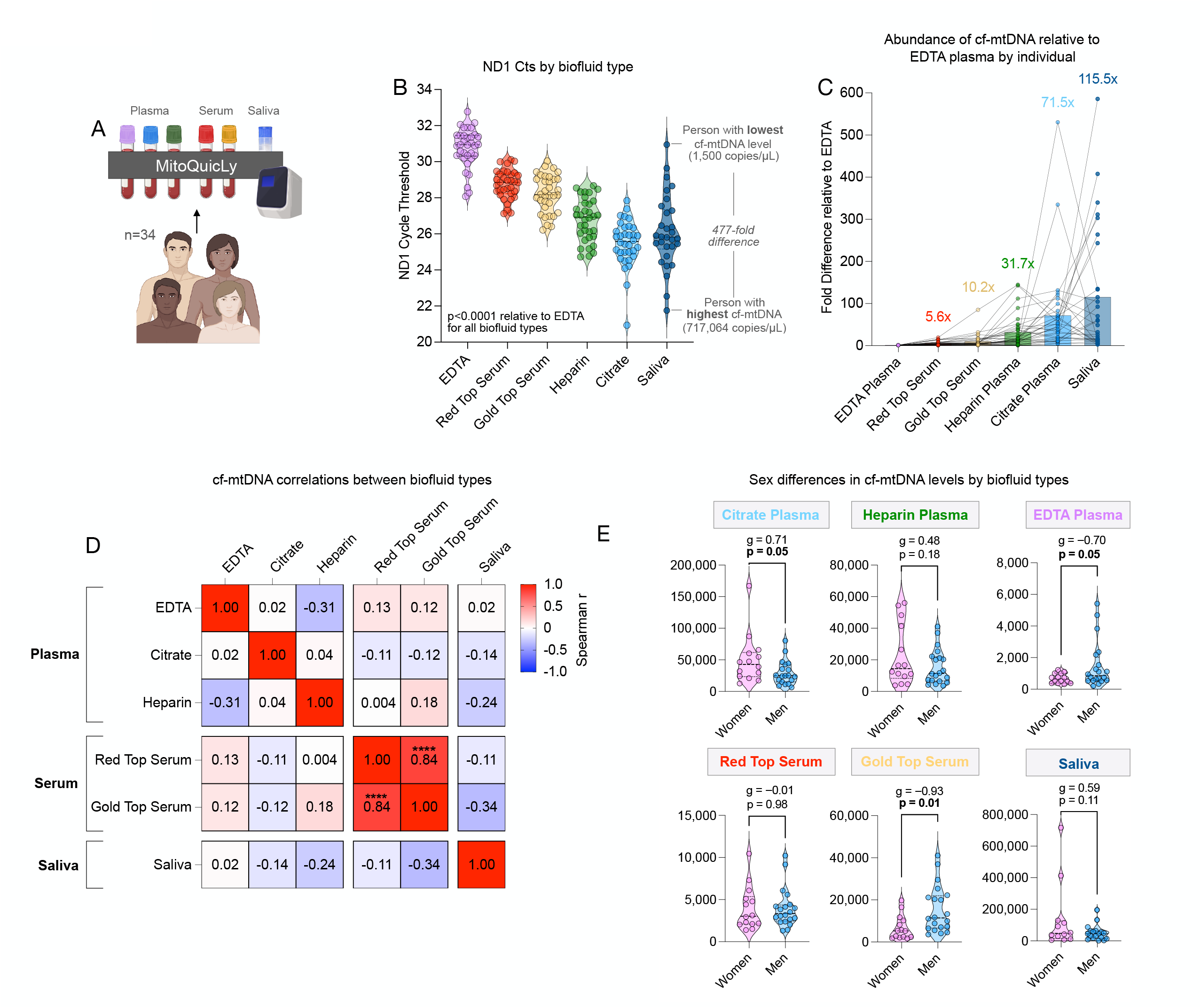
MitoQuicLy reveals sample type differences in cf-mtDNA. (**A**) Schematic of experimental design for MitoQuicLy cf-mtDNA analysis across 6 biofluids. (**B**) cf-mtDNA cycle threshold by sample type, revealing differences in cf-mtDNA levels between sample types, and substantial inter-individual differences. Each datapoint represents a participant (n=34 per biofluid). (**C**) Fold differences in cf-mtDNA copies/µL normalized to EDTA plasma cf-mtDNA copies/µL for each individual (each participant set to 1 for EDTA, connected by a line across sample types). The average fold difference for each sample type across the cohort is indicated. (**D**) Spearman’s r of cf-mtDNA copies/µL between sample types. Only red-top and gold-top serum cf-mtDNA levels show a significant correlation with each other. Plasma cf-mtDNA levels are not significantly correlated with serum or saliva, and saliva cf-mtDNA levels are not significantly correlated with any sample type. (**E**) Sex differences in cf-mtDNA levels by sample type. Shown are Hedges’ g (standardized effect size) and P values from unpaired t tests, n=34 participants (14 women, 20 men).

**Figure 7:**
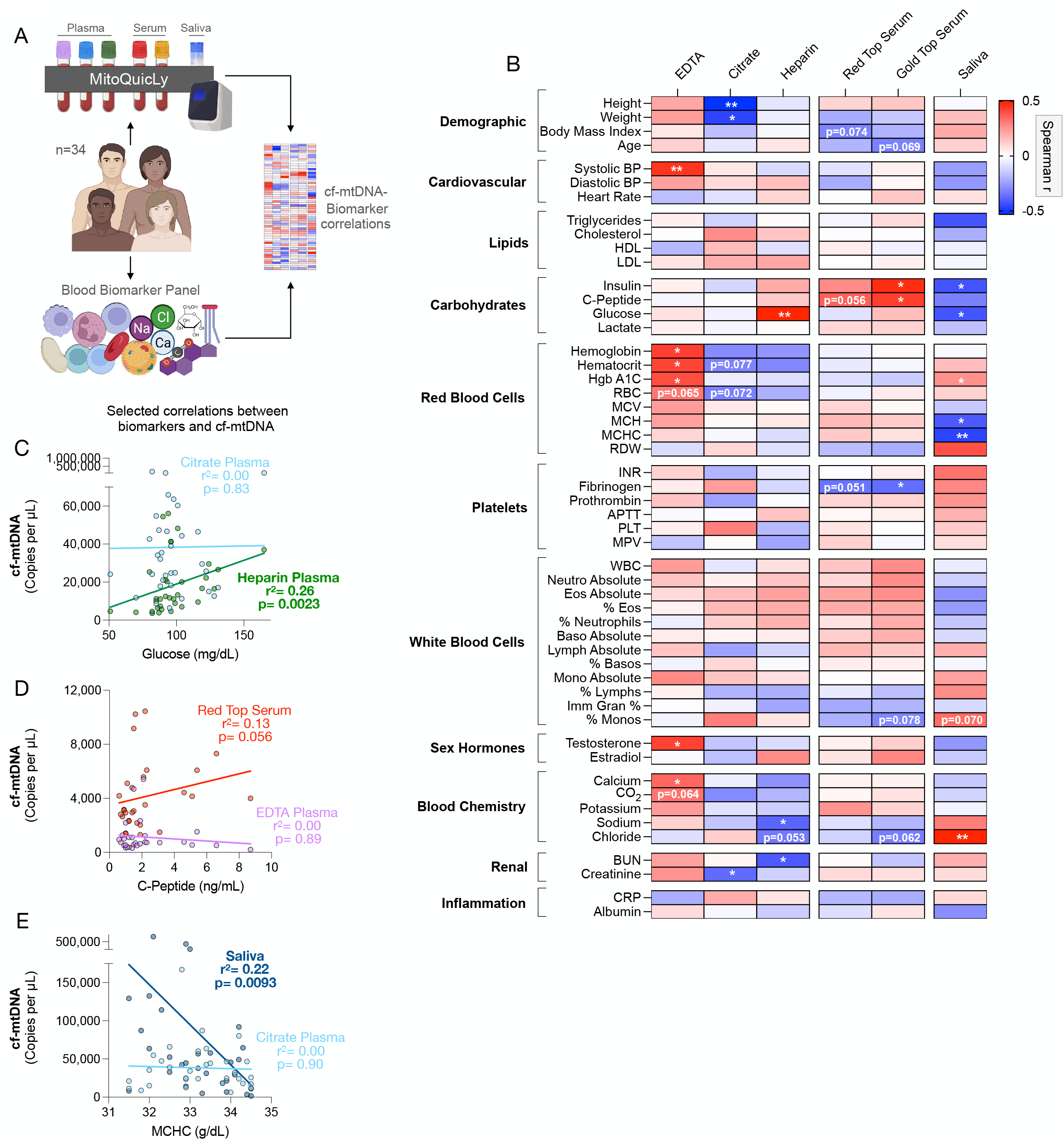
MitoQuicLy reveals different biological associations with cf-mtDNA by sample type. (**A**) Experimental design to combine MitoQuicLy cf-mtDNA levels from 6 biofluids and a standard blood biomarker panel. (**B**) Heatmap of correlations (Spearman’s r) between cf-mtDNA levels measured in each biofluid (n=34 individuals per sample type) and blood biomarkers. P values are not corrected for multiple comparisons. (**C-E**) Selected correlations (Spearman r) between blood biomarkers and cf-mtDNA levels. Each datapoint represents a participant (n=34; 14 women, 20 men). Spearman’s r correlations and p-values not corrected for multiple testing; p<0.05*, p<0.01**, p<0.001***, p<0.0001****. *Abbreviations*: *BP*, blood pressure; *HDL*, high-density lipoprotein; *LDL*, low-density lipoprotein; *Hgb A1C*, hemoglobin a1c; *RBC*, red blood cell count; *MCV*, mean corpuscular volume; *MCH*, mean corpuscular hemoglobin; *MCHC*, mean corpuscular hemoglobin concentration; *RDW*, red cell distribution width; *INR*, international normalized ratio of prothrombin time; *APTT*, activated partial thromboplastin time; *PLT*, platelet count; *MPV*, mean platelet volume; *WBC*, white blood cell count; *BUN*, blood urea nitrogen; *CRP*, c-reactive protein.

## 3. Results

### 3.1. Identifying failure points in plasma lysis and cf-mtDNA yield

MitoQuicLy is based on a lysis method optimized for single muscle cells (Murphy et al., 2008), which used a buffer composed of 50 mM Tris-HCl, 0.5% Tween, and 0.2 mg/mL Proteinase K. To extend this method to cf-mtDNA and cf-nDNA quantification in biofluids, we first sought to establish the proper concentration of the sample to be lysed. A series of lysis reactions of EDTA plasma (pooled from n=10 individuals) produced a stringy white precipitate in the 3 most concentrated dilutions but not in lower concentrations (Figure 1C). Using qPCR, we confirmed that relative to the supernatant, the precipitate contained concentrated DNA (95x more mtDNA than supernatant) (Figure 1D). B2M was not detected in the supernatant (Ct>40) but was detected in the precipitate, indicating that lysing EDTA plasma at high concentrations in this buffer causes DNA to precipitate out of solution.

Unprecipitated samples showed Cts that were inversely proportional to the plasma concentration (i.e., adding more plasma to the lysis reaction increases the number of DNA copies detected): slope=-1.14, r^2^=0.94, p=0.02 (Figure 1E). However, samples with precipitate (i.e. higher concentration samples) showed high Cts without proportionality: slope=-0.04, r^2^= 0.68, p=0.38. This indicated the existence of a “failure zone” of plasma concentration at which cf-mtDNA cannot accurately be quantified. To identify the concentration at which failure occurs, we lysed EDTA plasma from 10 individuals, each diluted at 28 different concentrations from 10% to 0.025% plasma (vol/vol) (Figure 1F). Across 10 individuals, failure clearly occurred at or above 2% plasma (Figure 1F, grey shading). Below 2% plasma, our lysis assay was linear to the 0.025% (1:4000) dilution.

### 3.2. Modifying buffer component concentrations improves DNA yield

This initial result suggested that plasma lysis could be performed at a 1% sample concentration. However, such low concentration did not yield reliable results (data not shown), which motivated a systematic optimization of each buffer component to improve the DNA yield. Therefore, we tested 10 concentrations of each of the three buffer components (Tris-HCl, Tween, Proteinase K) in citrate plasma and saliva from n=6 individuals (3 women, 3 men, ages 20-40; Supplemental Figure 2). Citrate plasma was used instead of EDTA plasma because pilot experiments showed citrate plasma had higher cf-mtDNA concentrations, which allowed for more accurate quantification. Compared to the original buffer used above, cf-mtDNA yield in plasma was improved by using either 114 mM Tris-HCl (+34% yield), 6% Tween (+68%), or 2 mg/mL Proteinase K (+36.5%). In saliva, the cf-mtDNA yield was improved by using either 114 mM Tris-HCl (+253% yield), 6% Tween (+32.5%), or 2 mg/mL Proteinase K (+53.7%). Thus, DNA yield was substantially improved by using higher concentrations of lysis buffer component.

Based on these results, we tested each combination of the original low (L) or high (H) concentration of each buffer component (Figure 2A). This experiment provided 5 potential buffers, each named for their concentration of each component (e.g., HHL = *High* Tris, *High* Tween, *Low* Proteinase K). We then tested each buffer simultaneously in citrate plasma and saliva from the same 6 individuals. Compared to the original LLL buffer in both plasma and saliva, all H buffers significantly increased cf-mtDNA yield 3-4-fold (p<0.0001) (Figure 2A-B). Comparing the median copy number obtained from each buffer showed that of all buffers tested, HHL consistently provided the highest cf-mtDNA yield (Figure 2D).

### 3.3. Titration of HHL buffer

We then deployed the HHL buffer in citrate plasma and saliva and performed a 1:2 serial dilution ranging from 20% to 0.078% sample (vol/vol) (Figure 3A and E). Unlike our original titration (Figure 1F), ND1 copies/µL scaled linearly with input biofluid in lysis reactions containing 0.078% to 10% plasma, and in all tested saliva reactions (0.078% to 20%, Figure 3B and F). By comparison, cf-nDNA was not detected in citrate plasma (Supplemental Figures 3A-3C) but was linearly detected in saliva (Supplemental Figures 3D-3F). Correcting for the dilution factors revealed the stability of cf-mtDNA yield across a wide range of sample inputs.

Accordingly, the rank-order of all participants was consistent across all concentrations, illustrating the stability of the assay across a broad range of biofluid concentrations (Figure 3C and G). Accordingly, excluding 20% plasma, there was excellent reliability among different lysis reactions (Spearman’s r > 0.9) for both plasma and saliva across all concentrations tested (Figure 3D and H). To select an input volume that would be sample-sparing while not too small (to minimize pipetting error and subsampling error (Taylor et al., 2019)), we opted to run MitoQuicLy at a 5% blood or saliva concentration.

### 3.4. Quantifying assay variability

To quantify the technical variability of the MitoQuicLy method, we pooled source biofluids from all participants (n=34) in our small cohort. We independently ran 96 lysis reactions from citrate plasma, EDTA plasma, red top serum, and saliva (Figure 4A). The coefficients of variation (CV) between lysis replicates were 8.7% for plasma, 13.3% for serum, and 8.2% for saliva (Figure 4B-E). EDTA plasma has much less cf-mtDNA than these other sample types, and the higher resulting cycle threshold values led to a 89.2% CV between replicates. Because there is much less cf-nDNA, higher CVs were seen in cf-nDNA levels from the same samples: 58.1% for citrate plasma, 50% for EDTA plasma, 27% for red top serum, and 16.7% for saliva (Supplemental Figure 4). Thus, for most biofluids except EDTA plasma, MitoQuicLy should detect sample-to-sample differences in cf-mtDNA concentrations of >10-15%. This sensitivity threshold lies below previously reported basal inter-individual differences in our data (see Figure 6) and by other laboratories (Lindqvist et al., 2016; Lindqvist et al., 2018; Nakahira et al., 2013; Rosa et al., 2020), and also below the magnitude of previously reported within-person changes in cf-mtDNA over time in blood and saliva (Cushen et al., 2020; Hummel et al., 2018; Trumpff et al., 2019; Trumpff et al., 2022). We detected no effect of well position on the 96-well lysis plate, ruling out sample location as a potential contributor to variability.

### 3.5. Comparison to silica membrane DNA isolation kit

Many laboratories use silica membrane-based DNA extraction kits (i.e., columns) to isolate DNA. To compare MitoQuicLy to the standard silica column approach, we isolated DNA using both column and MitoQuicLy on 50 serum and plasma samples (n=10 participants spanning the spectrum of inter-individual differences observed, 5 biofluid types per participant). The resulting dataset covers a wide range of cf-mtDNA concentrations spanning 2-3 orders of magnitude and therefore represents a robust technical comparison.

Overall, there was high agreement (r^2^=0.74, p<0.0001) between both DNA extraction methods (Figure 5A). One notable finding is that the final extracted DNA from MitoQuicLy (lysed biofluid) and column (purified DNA) produced similar ND1 Cts (Figure 5A). However, the biofluid sample input for the column was 100 µL, compared to 10 µL of the same biofluid for MitoQuicLy, pointing to an approximately 10x higher absolute DNA yield with MitoQuicLy. The difference in yield is likely attributable to the well-known retention of DNA in silica-based columns, which can also vary between the smaller mitochondrial and larger nuclear genomes (Guo et al., 2009). Another notable observation from these results is that ND1 Cts clustered by biofluid sample type (Figure 5B), pointing to large potential differences in cf-mtDNA abundance by sample type corroborated by both the column and lysis methods.

The comparative advantages of MitoQuicLy are summarized in Figure 5C. MitoQuicLy is higher throughput and can accommodate 96 samples in a plate without adding much time burden, while each additional sample of column isolation requires additional pipetting time. Finally, standard column-based DNA extraction costs □$3.00 USD per sample for a large number of samples, while MitoQuicLy costs ∼$0.20, at least an order of magnitude less expensive.

### 3.6. Biofluid differences in cf-mtDNA concentrations

To systematically examine differences in cf-mtDNA concentrations between biofluids, we deployed MitoQuicLy in all participants from Cohort 2 to compare cf-mtDNA levels in 6 concurrently collected biofluid sample types (total n=200 samples): three plasma samples (*EDTA, Citrate*, and *Heparin*), two serum samples (*Red top*, with clot activator; *Gold top*, with clot activator + gel separator), and one *saliva* sample (Salivette) (Figure 6A). Significant differences in cf-mtDNA levels between sample types were observed (Hedges’ g=2.09-4.36, ps<0.0001, Figure 6B). Of all samples tested, EDTA plasma had the lowest cf-mtDNA levels, while citrate plasma and saliva had the highest.

Interestingly, the inter-individual differences in cf-mtDNA levels varied depending on the choice of biofluid sample type. Figure 6C shows the average fold difference in cf-mtDNA levels from the same participant compared to their EDTA plasma value (each line is a participant). Compared to EDTA plasma cf-mtDNA levels, the fold differences were 5.6-10.2 for serums, 31.7-71.5 for heparin and citrate plasma, and 115.5 for saliva. Thus, cf-mtDNA levels vary by over two orders of magnitude between biofluid types.

One potential reason for these large biological differences could be that the chemistry of the lysis process operates differently in different biofluid types, yielding incomplete lysis or DNA quantification in some sample types. However, similar albeit smaller differences between biofluid types were also observed when DNA was extracted with the column-based method (Supplemental Figure 5A). Consistent with high agreement across column and lysis methods (Figure 5B), across the 5 biofluid types quantified by the MitoQuicLy and column-based method, the rank order of cf-mtDNA levels between individuals was comparable, with the exception of heparin plasma (Supplemental Figure 5A & B). Heparin is a known inhibitor of qPCR reactions (Meddeb et al., 2019b). In the column method, relatively little heparin is likely to contaminate the qPCR reaction, while in MitoQuicLy, there is no additional purification to remove heparin from the sample, which could cause DNA levels in heparin plasma to appear lower from samples isolated via MitoQuicLy compared to those isolated with the column method. Evidence of contaminating heparin interfering with qPCR can be seen in Supplemental Figures 5C-D. cf-nDNA was detected in heparin plasma prepared with the column method (Figure S5C), while it was not detected in heparin plasma prepared via MitoQuicLy (Figure S5D).

### 3.7. cf-mtDNA across biofluids, sex, and common biomarkers

We then examined the correlation in cf-mtDNA levels between biofluids. Within each participant, cf-mtDNA levels between biofluids were not correlated (Figure 6D). This means, for example, that relative to the group, the participant with the highest cf-mtDNA levels in heparin plasma may not have the highest cf-mtDNA level in other biofluid types (see Figure 6C). The cf-mtDNA levels in blood-derived plasma types and saliva were also independent, pointing to inherent biological differences between biofluid types. Only the two serum types were significantly correlated (Spearman’s r=0.84, p<0.0001).

As a proof-of-concept analysis to examine how the lack of agreement among cf-mtDNA levels across sample types could influence conclusions for targeted research questions, we first evaluated sex differences in cf-mtDNA levels. Prior research has shown either no difference in cf-mtDNA levels by sex (Jylhävä et al., 2014; Meddeb et al., 2019a; Zhong et al., 2007), or higher levels in men than in women (Kananen et al., 2022; Yuwono et al., 2021). In our sample, the effect sizes between women and men varied depending on the biofluid used (Figure 6E). In gold top serum, cf-mtDNA was higher in men than women (g=-0.93), consistent with our previous findings in serum from an independent cohort (Trumpff et al., 2019), while citrate plasma cf-mtDNA showed the opposite trend (g=0.71). In this small sample, red top serum cf-mtDNA did not vary by sex (g=-0.01). Together, this suggested that the choice of biofluid type may have substantial analytical implications for simple research questions.

To examine this point more systematically in relation to commonly used clinical variables, we computed correlation coefficients of cf-mtDNA levels in each biofluid type with demographic and cardiovascular measurements, together with blood-based chemistry, hematological, metabolic, and inflammatory biomarkers (Figure 6B). Consistent with the lack of correlation in cf-mtDNA between biofluid types, each biofluid type exhibited a unique correlation profile with blood biomarkers. Although both serum samples’ cf-mtDNA levels were positively correlated with insulin and C-peptide concentrations, EDTA plasma cf-mtDNA was correlated with hemoglobin, hematocrit, and glycated hemoglobin (Hgb A1c). Selected correlations divergent across biofluid types are shown in Figure 6C-E. While this small study should not be viewed as providing definitive estimates of the correlation between cf-mtDNA and blood biomarkers, these data illustrate important biological divergences between biofluid types, and explain why previous findings using different biofluid types may have yielded conflicting results. This systematic comparison provides an empirical foundation for additional, well-powered studies examining the nature and physiological significance of cf-mtDNA in different biofluids.

## 4. Discussion

We have developed MitoQuicLy, a lysis-based method to measure cf-mtDNA in human plasma, serum, and saliva. This method uses a small input volume and relatively inexpensive reagents and is scalable for high-throughput sample processing. MitoQuicLy can be used by researchers to sensitively detect and quantify cf-mtDNA in copies/µL in a variety of biofluid sample types and is easily scalable. Our comparative results across biofluid types highlight the need for additional research to decipher the biological nature of cf-mtDNA in human blood and saliva, which can then inform the selection of the ideal biofluid type for specific clinical and research applications.

One major new insight gained from this study is the differences in cf-mtDNA levels between biofluid types, even when concurrently collected from the same individuals. This observation is consistent with differences previously observed between serum versus plasma cf-mtDNA levels, which have been attributed to the release of mtDNA from activated platelets (Boudreau et al., 2014; Rosa et al., 2020; Xia et al., 2009). Other studies also have found elevated levels of cf-nDNA or circulating tumor DNA in serum compared to plasma (for a review, see (El Messaoudi et al., 2013)). These sample type differences indicate potentially divergent biology in the origin of cf-mtDNA. In particular, the large differences and lack of correlation in cf-mtDNA levels from EDTA, citrate, and heparin plasma suggests that cf-mtDNA in each type of plasma could be derived from different secretion mechanisms, cell-types, or type of pools of extracellular vesicles. This interpretation is supported by the unique correlation patterns between plasma cf-mtDNA levels and blood-based biomarkers. These data highlight the need for future research to understand the differences in the biology of these sample types and whether there are appropriate ways to compare them. Serum cf-mtDNA levels shared ∼70% of their variance (r=0.84) and showed similar correlation patterns with blood biomarkers, suggesting that findings from gold or red top tubes may be comparable, even if not identical. One factor contribute to this difference may be the presence of a “separation gel” in gold-top tubes, which may affect the final composition of the serum compared to the red-top serum tube. Finally, saliva cf-mtDNA levels were not associated with either plasma or serum cf-mtDNA levels, indicating that salivary cf-mtDNA may not reflect circulating blood cf-mtDNA levels and instead represent different cf-mtDNA biology deserving further research.

One question that naturally arises from our data is: “How do you know which tube to use?”. To that end, we are reminded that different clinical biomarkers have acceptable and unacceptable biofluid types. For example, most platelet function assays require blood drawn in citrate tubes (to prevent platelet activation). Lipid levels are typically measured in serum, not plasma. And catecholamines are quantified in heparin plasma. Our data and that of others unambiguously indicate that cf-mtDNA levels across biofluid types are not equivalent. Again, this highlights the need for additional knowledge about the origin and nature of these differences. Clarifying the underlying biology of cf-mtDNA across biofluid types would help establish two main points. First, it is possible that one tube type is indeed “optimal” and should become the gold standard. Alternatively, different tube types may allow quantifying different “subtypes” of cf-mtDNA (e.g., enclosed in medium vs. large vesicles, whole mitochondria, naked DNA, etc.), and therefore have distinct biological significance. If this was the case, there may not be an optimal universal tube type. Instead, different tube types could allow investigators to discern different aspects of cf-mtDNA biology more or less relevant to the specific research question via the selection of particular tube types.

MitoQuicLy has several notable advantages over alternative approaches. The *first* is throughput. By performing plasma lysis directly in a 96-well plate format, compared to DNA extraction procedures that are carried out individually in spin columns, the additional time required per additional sample is relatively minimal. Additionally, after lysis is complete, a multichannel pipette or automated liquid handling station can efficiently transfer lysates into a qPCR plate for analysis. Thus, MitoQuicLy is relatively easily scalable to thousands of samples, which is advantageous for both repeated-measures study designs and large-scale epidemiological studies involving hundreds or thousands of samples. A *second* advantage is the lack of extraction bias. Unlike with silica-based membranes (Guo et al., 2009), the single-step lysis method avoids selective retention/loss of mitochondrial over the nuclear genome. As a result, the cf-DNA concentration in the biofluid after qPCR can be calculated directly by correcting for the 1:10 dilution factor. The *third* advantage is the use of small input volumes. As noted above, the 10 µL volume requirement economically uses limited human biological samples. The *fourth* advantage is cost. At an approximate cost of USD$0.20/sample, our method is significantly cheaper than other DNA isolation methods, which can become considerable when analyzing large sample sizes. *Finally*, it is also well known that “science has a plastics problem” (Urbina et al., 2015) compounded by the use of disposable plastic consumables at multiple steps of molecular biology techniques. Compared to silica-based membrane kits and other multi-step methods, the single-step lysis approach in one (two if run in duplicates) standard microplate for 96 samples minimizes the mass of single-use plastic per sample.

The MitoQuicLy method also has several limitations. *First*, the 200 µL lysis reaction carried out at 5% plasma concentration can only accommodate a small plasma volume (10 µL). While this is an advantage in research or clinical context where sample volume is limited, the small volume pipetted can increase variability when the cf-mtDNA concentration is low. In this case, variability is due to subsampling error, in which different aspirations of the same source biofluid sample may have different concentrations of mtDNA than the sample as a whole (Taylor et al., 2019). *Second*, because the sample is diluted, a relatively small number of cf-mtDNA molecules are loaded into each qPCR well. Therefore, for lower abundance samples (i.e., EDTA plasma), this yields relatively high (∼30) Ct values, which inherently have higher technical variability due to stochastic amplification and measurement uncertainty (Taylor et al., 2019). Although the variability of our measurement of EDTA cf-mtDNA levels is relatively high, the values are highly correlated with those obtained from column kit extraction (Figure 5B and Supplemental File 2: r^2^=0.70, p=0.0024), suggesting the results obtained from MitoQuicLy are comparable to those from other techniques. Due to the known variability of our method, we recommend lysing each biofluid sample twice (duplicates), in addition to the standard three technical replicates per qPCR reaction in accordance with the MIQE guidelines (Bustin et al., 2009). Across hundreds of plasma, serum, and saliva samples run in different batches on separate days, the average technical variation between two MitoQuicLy runs on the same sample is 11.3% (Red top serum), 14.78% (EDTA plasma), and 14.2% (saliva), highlighting the stability and also the detection limit of the optimized assay. *Finally*, because qPCR is performed directly on the lysate without additional purification steps, the isolated DNA is not sufficiently pure for other applications, such as sequencing. If needed for other applications than cf-mtDNA quantification, the DNA from the MitoQuicLy lysate can be purified using standard methods (e.g., silica membrane, magnetic bead-based isolation).

Overall, we report a simple, high-throughput, cost-effective cf-mtDNA extraction method that can be deployed at scale to quantify cf-DNA concentrations in a variety of human biofluids. The initial optimization steps and systematic MitoQuicLy analyses across six sample types identified large and significant differences in cf-mtDNA levels between different commonly used biofluids. Moreover, cf-mtDNA levels in each biofluid showed unique correlation patterns with blood biomarkers, which may explain previous discrepancies in the literature and points to potentially divergent and unresolved cf-mtDNA biology between plasma, serum, and saliva.

Further studies in large cohorts are needed to deepen these observations and further define the biological relevance and clinical correlates of cf-mtDNA levels. The MitoQuicLy protocol (Supplemental File 1) is a scalable approach to assess cf-DNA levels in a diversity of biofluids.

## Supporting information

Supplemental File 1: MitoQuicLy Protocol

Supplemental File 2: MitoQuicLy Data and Figures

## Acknowledgements

We are grateful to the participants for this study and to the Center for Advanced Laboratory Medicine (CALM) Laboratory for the measurements of blood biomarkers.

## Supplemental Material

This article contains 5 supplementary figures and 2 supplemental files.

## Supplemental Figure Legends

**Figure S1:**
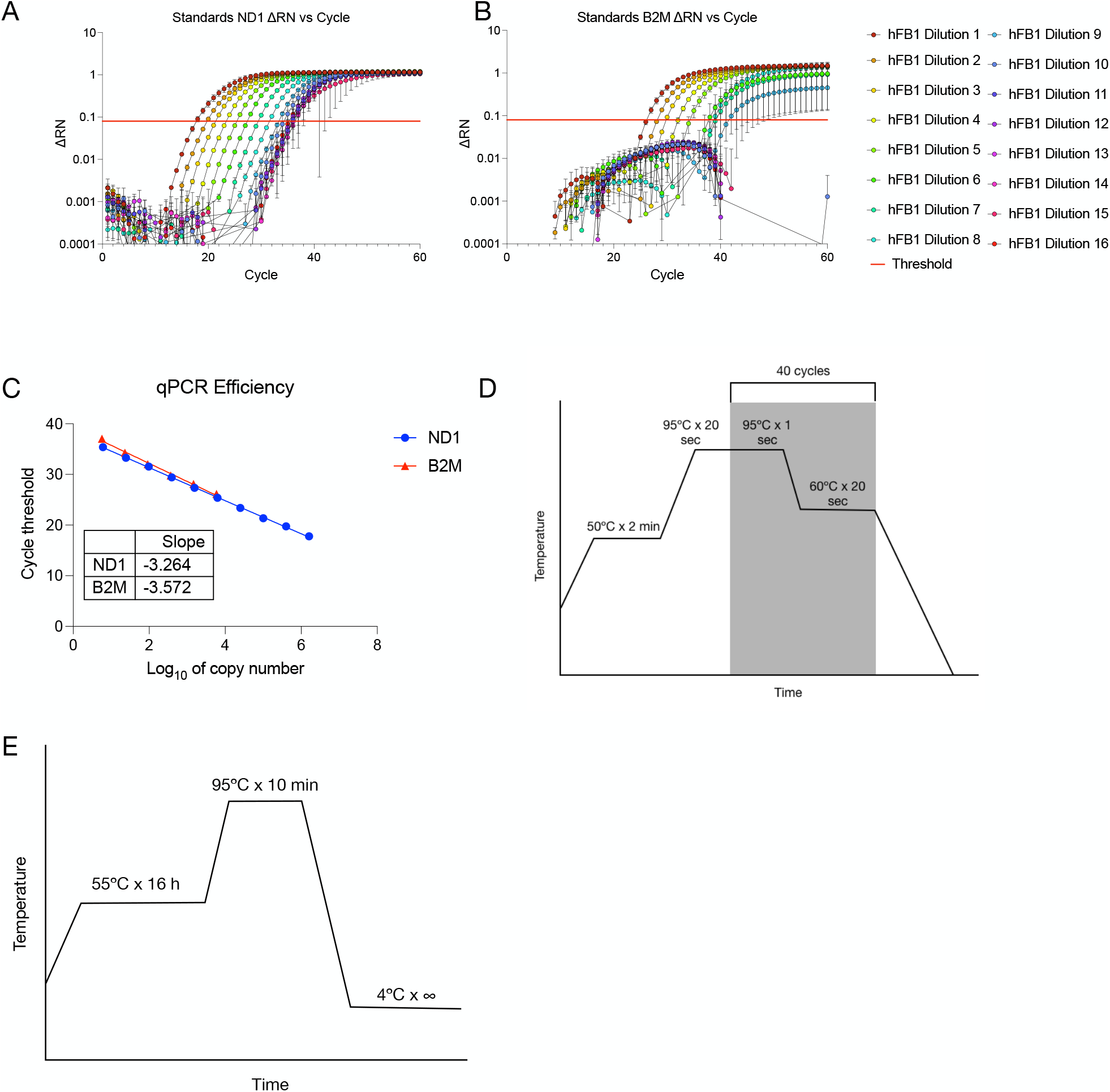
qPCR validation. (**A and B**) hFB1 standard curves. Primary human fibroblasts were collected and DNA was isolated using the lysis method and diluted to a concentration of 1 µg/mL. 16 serial 1:4 dilutions were performed from this stock. mtDNA and nDNA was quantified using ND1-B2M duplex qPCR. A ΔRN threshold of 0.08 was used to determine cycle thresholds (red line). (**C**) Efficiency of ND1-B2M qPCR. (**D**) qPCR cycling conditions. (**E**) MitoQuicLy incubation times and temperatures.

**Figure S2:**
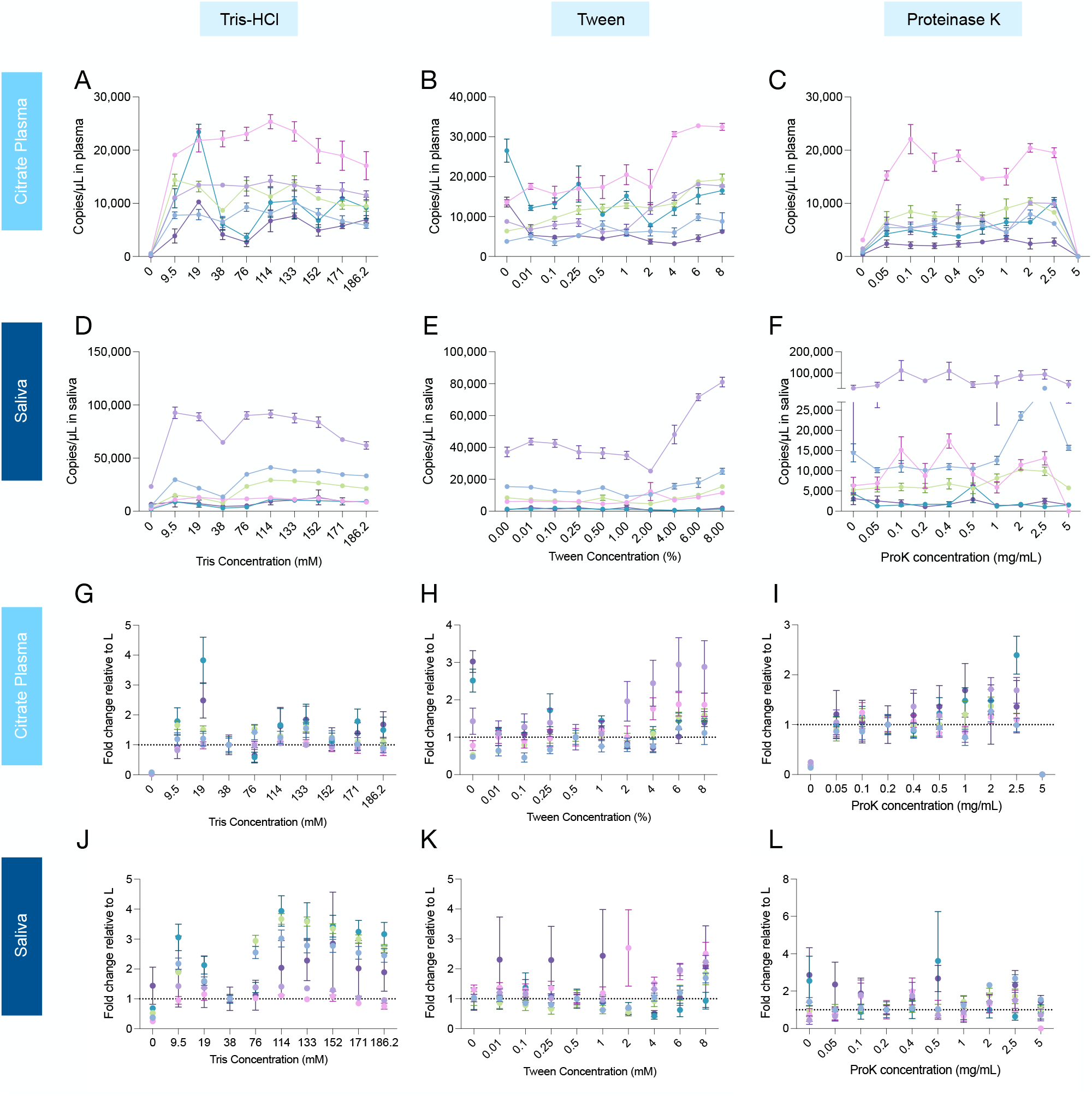
Testing buffer component concentrations. Ten concentrations of each buffer component were tested in both citrate plasma (**A-C**) and saliva (**D-F**). qPCR was performed and mtDNA copies/µL were computed for each sample. (**G-L**) ND1 copies/µL at each buffer concentration were normalized to the original concentration used to identify which concentrations provided substantially improved yield. For Tris-HCl, 114 mM was optimal. For Tween, 6% was optimal. For ProK, 2.5 mg/mL improved yield, but 5 mg/mL caused qPCR to fail.

**Figure S3:**
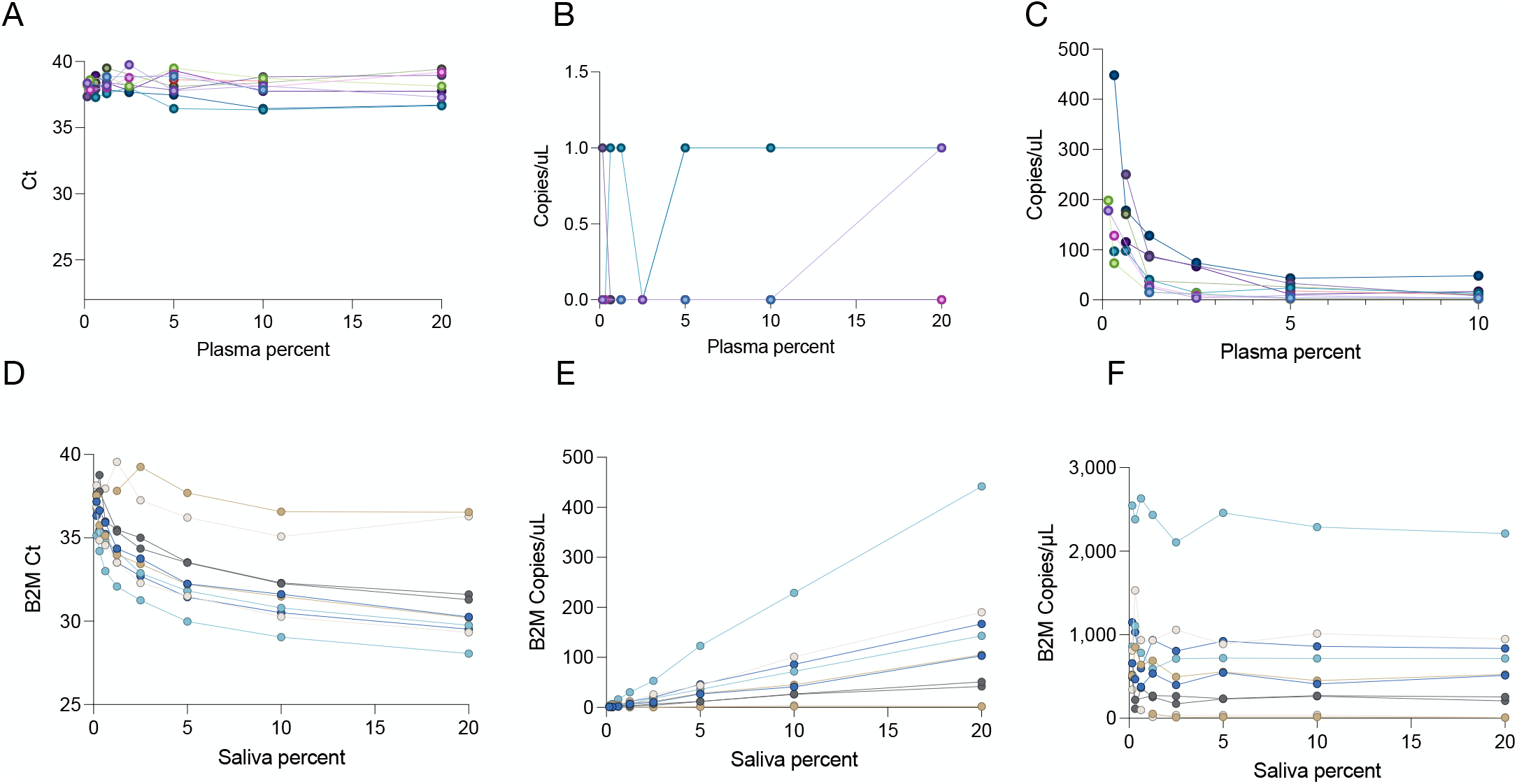
Titration of sample concentration: nDNA. Since HHL significantly improved yield, we asked whether it could handle an increased concentration of plasma or saliva. Citrate plasma and saliva from n=10 people was lysed at 8 2-fold serial dilution points from 20% to 0.078% sample (vol/vol). qPCR was performed for the B2M (nuclear) gene. Each person’s serial dilution is represented by points on a connected line. (**A and D**) B2M cycle threshold at each dilution point. (**B and E**) Copies/µL computed from cycle threshold by correlation with a standard curve of known concentration. In citrate plasma, cf-nDNA was at or near the limit of detection at all concentrations tested. (**C and F**) Dilution corrected copies/µL in plasma and saliva. From 0-20% saliva, we can reliably rank-order the samples. n=10 individuals, 5 women, 5 men.

**Figure S4:**
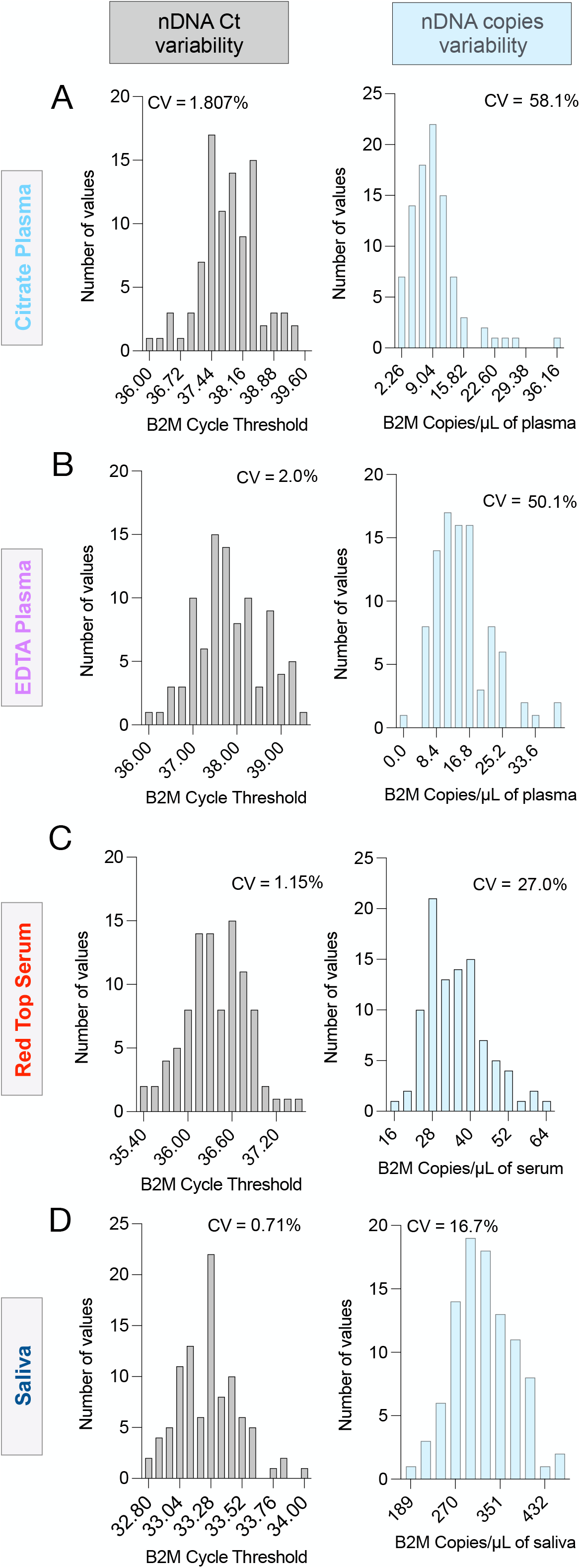
Assay variability. 96 replicates each of pooled sample were analyzed using MitoQuicLy. Histograms of citrate plasma (**A**), EDTA plasma (**B**), serum (**C**), and saliva (**D**) nDNA levels are shown in terms of Ct and copies/µL. n=34 people (14 women, 20 men).

**Figure S5:**
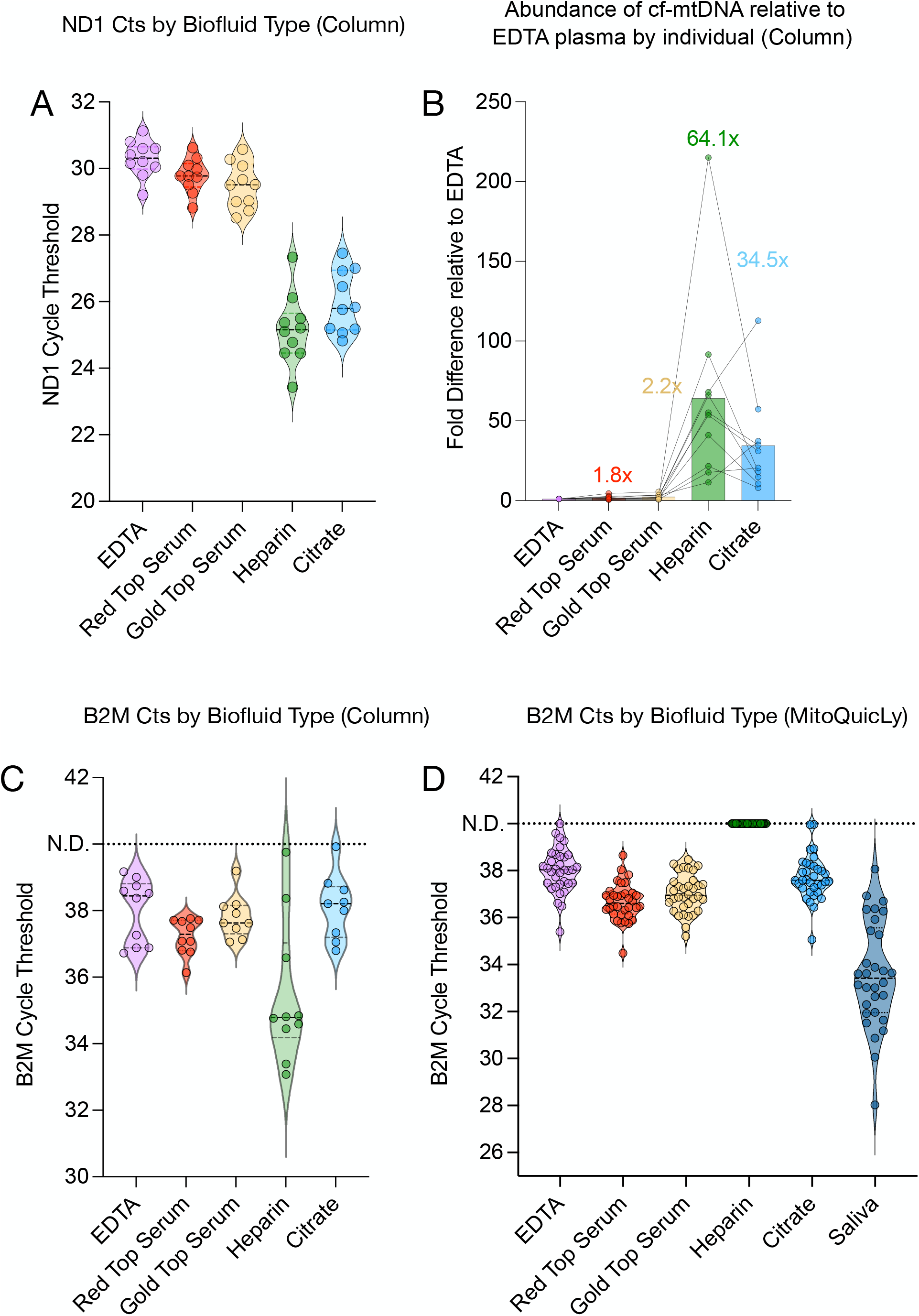
Sample type differences in cf-mtDNA. (**A**) cf-mtDNA cycle threshold by sample type after isolation using Column method. Citrate and Heparin have significantly more cf-mtDNA than EDTA, Red Top Serum, or Gold Top Serum (p<0.001), but no significant difference was seen between EDTA/Red Top Serum/Gold Top Serum or between Citrate/Heparin. (**B**) Intra-individual fold differences in cf-nDNA copies/µL normalized to EDTA plasma cf-nDNA copies/µL. Individual participants’ cf-nDNA levels are connected by a line. The average fold increase for each sample type is shown above. (**C**) B2M Cycle threshold in 10 participants cell-free plasma or serum isolated via column method (**D**) B2M Cycle threshold in 34 participants’ cell-free plasma, serum, or saliva isolated using MitoQuicLy. For A, B, and C, n=10 participants (5 women, 5 men). For D, n=34 participants (14 women, 20 men).

**Supplemental File 1. Detailed point-by-point MitoQuicLy Protocol**.

**Supplemental File 2. Source data, analyses and graphs (GraphPad Prism file)**.

