## Supplemental File 1: MitoQuicLy Protocol for "MitoQuicLy: a high-throughput method for quantifying cell-free DNA from human plasma, serum, and saliva"

**MitoQuicLy: Mitochondrial DNA Quantification in cell-free samples by Lysis**


### Summary:

This protocol describes procedures for collecting, isolating, and quantifying cell-free mitochondrial DNA (cf-mtDNA) and nuclear DNA (cf-nDNA) in humans using quantitative polymerase chain reaction (qPCR).


### Graphical summary:

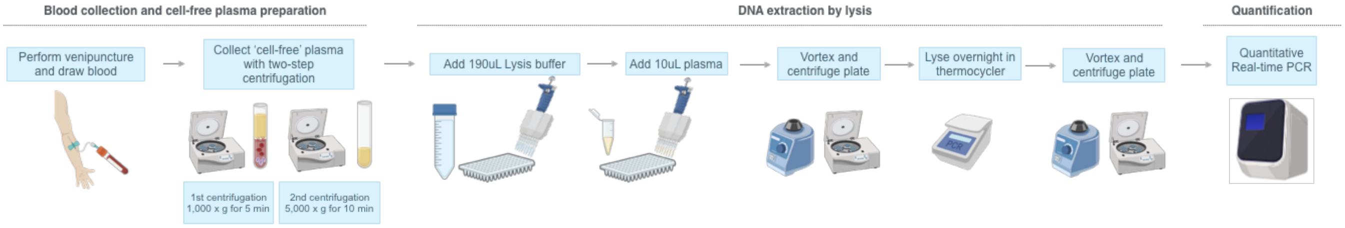


### Consumables, with catalog numbers:

BD Vacutainer Tubes (BD #363083)

Tris HCl (Sigma #T3253)

Tween 20, 10% (Sigma #P1379)

Nuclease free water (Thermofisher Scientific #AM9939)

Proteinase K (20mg/ml) (Thermofisher Scientific #AM2548)

Stericup Quick Release-GV Sterile Vacuum Filtration System – 500 mL (Millipore Sigma S2GVU05RE)

BrandTech 96-well semi-skirted plate (BrandTech #781375/VWR #10141-434)

BrandTech 8-strip domed tube caps (BrandTech #781340/ VWR #80087-132)

2x TaqMan Universal MasterMix Fast (LifeTech #4444965)

MicroAmp™ Optical 384-Well Reaction Plate with Barcode (ThermoFisher #4309849)

MicroAmp™ Optical Adhesive Film (ThermoFisher #4311971)

Validated mtDNA/nDNA Primer & Probe sequences (idtDNA.com)

Additional consumables not listed here: 1.5mL Eppendorf tubes, pipette tips as needed.

### Protocol to collect samples:

#### *Blood collection & cell-free plasma preparation*

- Perform venipuncture with standard butterfly needle or catheter (20 gauge or larger).
- Collect the first tube of 2-3mL of blood, which may contain contaminating genomic material from cellular damage during venipuncture, and use for other analytes or discard.

- Draw required volume of blood for cf-DNA measurements (ideally 5mL or more)[1] into BD Vacutainer tubes. We use Citrate tubes (BD #363083).
- Gently invert 10 times to ensure homogeneous mixing of anticoagulant with the blood sample.

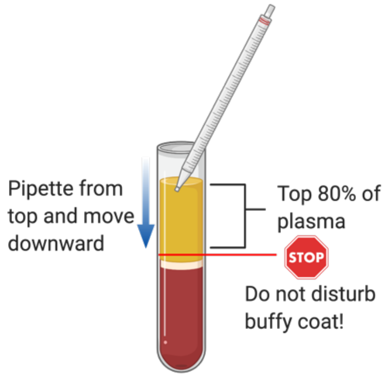


- Immediately (within 2 minutes, if possible) centrifuge blood tubes at 1,000g x 5 minutes at room temperature.
- Blood will separate into three layers, bottom layer is red blood cells, top layer is plasma, and the interface is the buffy coat. Using a serological pipette, slowly aspirate 80% of the plasma from the top of the tube without approaching the buffy coat.
- Transfer plasma to a new tube.
- Centrifuge plasma at 5,000g x 10 min at 4ºC to pellet remaining potential contaminating platelets.
- Slowly aspirate 80% of the plasma from the top of the tube without approaching the bottom of the tube. Transfer this supernatant – "clean plasma" – to a new tube.
- Mix gently with a pipette or invert to ensure that the clean plasma is homogenous.
- Aliquot into cryovials and store at -80ºC or proceed to DNA extraction by lysis.

### Lysis buffer for cf-DNA measurement

*In saliva, plasma, and serum*
*Keep everything sterile The below calculations provide sufficient lysis buffer for 1L of complete lysis buffer, or about 5,000 reactions*

**Tris-HCl (0.38M, pH 8.5)** (Sigma #T3253)

- 18.168 g of Tris HCl (MW = 157.60 g/mol) in 200ml of nuclease-free $dH_2O$
- pH to 8.5 with 5M NaOH (about 12.5mL, but measure pH continuously)
- Note: Accurate pH is critical for proper extraction. Make sure your pH meter has been recently calibrated.
- Make up to 300ml with nuclease-free $dH_2O$
- Sterile filter Tris-HCl buffer using a SteriCup

*(Make 6ml or 12mL aliquots, depending on if you want to run 1 or 2 plates at once)*

[1] While the final measurement of cf-mtDNA only requires about 20 µL of final product, we suggest 5mLs as a minimum blood volume because less volume makes the pipetting steps more technically difficult.

**Tween 20, 10%** (Sigma #P1379)

- 540ml of ultrapure $dH_2O$
- 60ml of Tween 20 (wash tip thoroughly or leave in solution)
- Sterile filter Tween 10% using a SteriCup

*(Make 12ml or 24mL aliquots, depending on if you want to run 1 or 2 plates at once)*

NB: Thawing these 10% Tween aliquots can be slow. I recommend using a 37ºC bead bath.

**$ddH_2O$ (Sterile, nuclease-free)** (Thermofisher Scientific #AM9939)

*(Make 5ml aliquots or use direct from bottle)*

**Proteinase K (20mg/ml)** (Thermofisher Scientific #AM2548)

- To be put in fresh from 20mg/ml (1:100 dilution, final 200ug/ml)

Store aliquots at -30C

**Recipe for 1ml Lysis Buffer**

300µl – Tris HCl
600µl – Tween 20
90µl – nuclease free $H_2O$
10µl – Proteinase K

| Examples of Lysis Buffer Calculations (including a bit of excess) | | | |
|---|---|---|---|
| **Component** | **48 Samples** | **96 Samples (1 plate)** | **2 Plates** |
| **Tris** | 3 mL | 6 mL | 12 mL |
| **Tween** | 6 mL | 12 mL | 24 mL |
| **Proteinase K** | 100 µL | 200 µL | 400 µL |
| **Nuclease-free water** | 900 µL | 1800 µL | 3.6 mL |
| **Total Volume** | 10 mL | 20 mL | 40 mL |

***Lysis***

Note: DNA molecules are often contained within lipid-based structures and packaged with DNA, which prevents their accurate quantification. Therefore, it is necessary to extract DNA prior to quantification. Numerous DNA extraction methods are available, but many rely on silica-based filter systems (e.g., DNeasy) which limits yield and biases nDNA and mtDNA ratios (Fazzini et al., 2018; Guo, Jiang, Bhasin, Khan, & Swerdlow, 2009; Picard et al., 2012). A high-throughput method with excellent yield has recently been described (Ware et al., 2020). Here we describe a simple method that relies on digestion via a lysis buffer.[2]

- Thaw all reagents and prepare lysis buffer as described above.
- This assay should be run in technical duplicates, so prepare two identical plates using the same samples in the same arrangement.
- Add 190 μL of lysis buffer in each well of a 96-well semi-skirted plate. We use BrandTech CAT#781375. Supplied in US by VWR CAT#10141-434.
- Add 10 μL of sample into each well.
- We recommend one extra well with 200 μL lysis buffer only to use as a "no template control".
- Seal wells with 8-strip domed caps. We use BrandTech #781340. Supplied in US by VWR #80087-132.
- To ensure proper mixing of the sample and buffer, vortex vigorously.
- To pull the plasma/buffer mixture to the bottom of the wells, quickly centrifuge at 1,000g for 10 sec before lysis.
- Proceed to heat-activated lysis of the samples in a Thermocycler for 16 hours at 55°C, followed by heat inactivation for 10 minutes at 95°C.

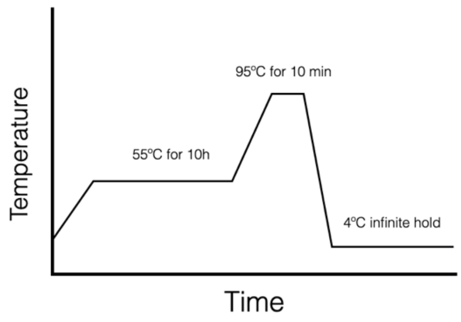


  - This digested sample can be used directly as template DNA in qPCR for cf-DNA measurements.
- Before performing qPCR, vortex and centrifuge again the template DNA-containing tube(s)/plate(s) at 1,000g for 10 sec at room temperature.
  - If not using within 24 - 48h, freeze the DNA samples at -80 °C or colder.

#### *qPCR plate preparation*

Note: Methods and reagents in this section may vary depending on the qPCR equipment used and the target amplicons. The following protocol is used for 20ul reactions:

- Samples are assayed in triplicate, so calculate 3x the total number of samples to be assayed. This provides the number of qPCR wells required.

[2] *Note*: The lysis buffer is composed of Tris-HCl, Tween 20, Proteinase K, and nuclease-free water. Tris-HCl stabilizes the pH of the buffer and protects nucleic acids from degradation. Tween 20 is a detergent used to solubilize lipid-rich membranes. Proteinase K is an enzyme used to break down proteins. Nuclease-free water is added to achieve optimal concentrations of each of these components.

- Prepare mastermix as described in the reagents sections in Appendix A to assay that number of wells, adding about 10-25% of excess volume to reduce risk of pipetting error (i.e, for 96 qPCR wells, prepare ~110 wells worth of mastermix) .
- Either manually or with an automated liquid handler, dispense 12 µL of the qPCR mastermix into each well of the qPCR plate.
- In pink below, see the arrangement of samples on a 96-well plate. In grey below, the arrangement of qPCR triplicates of both samples and standards on a 384-well plate.

| cf-mtDNA Plate 1 | 1 | 2 | 3 | 4 | 5 | 6 | 7 | 8 | 9 | 10 | 11 | 12 |
|---|---|---|---|---|---|---|---|---|---|---|---|---|
| A | 1 | 9 | 17 | 25 | 33 | 41 | 49 | 57 | 65 | 73 | 81 | 89 |
| B | 2 | 10 | 18 | 26 | 34 | 42 | 50 | 58 | 66 | 74 | 82 | 90 |
| C | 3 | 11 | 19 | 27 | 35 | 43 | 51 | 59 | 67 | 75 | 83 | 91 |
| D | 4 | 12 | 20 | 28 | 36 | 44 | 52 | 60 | 68 | 76 | 84 | 92 |
| E | 5 | 13 | 21 | 29 | 37 | 45 | 53 | 61 | 69 | 77 | 85 | 93 |
| F | 6 | 14 | 22 | 30 | 38 | 46 | 54 | 62 | 70 | 78 | 86 | 94 |
| G | 7 | 15 | 23 | 31 | 39 | 47 | 55 | 63 | 71 | 79 | 87 | 95 |
| H | 8 | 16 | 24 | 32 | 40 | 48 | 56 | 64 | 72 | 80 | 88 | 96 |

| | 1 | 2 | 3 | 4 | 5 | 6 | 7 | 8 | 9 | 10 | 11 | 12 | 13 | 14 | 15 | 16 | 17 | 18 | 19 | 20 | 21 | 22 | 23 | 24 |
|---|---|---|---|---|---|---|---|---|---|---|---|---|---|---|---|---|---|---|---|---|---|---|---|---|
| A | 1 | 1 | 1 | 9 | 9 | 9 | 17 | 17 | 17 | 25 | 25 | 25 | 33 | 33 | 33 | 41 | 41 | 41 | hfB1 Standard 1 | hfB1 Standard 1 | hfB1 Standard 1 | | | |
| B | 49 | 49 | 49 | 57 | 57 | 57 | 65 | 65 | 65 | 73 | 73 | 73 | 81 | 81 | 81 | 89 | 89 | 89 | hFB13 Standard | hFB13 Standard | hFB13 Standard | | | |
| C | 2 | 2 | 2 | 10 | 10 | 10 | 18 | 18 | 18 | 26 | 26 | 26 | 34 | 34 | 34 | 42 | 42 | 42 | hFB1 Standard 2 | hFB1 Standard 2 | hFB1 Standard 2 | | | |
| D | 50 | 50 | 50 | 58 | 58 | 58 | 66 | 66 | 66 | 74 | 74 | 74 | 82 | 82 | 82 | 90 | 90 | 90 | | | | | | |
| E | 3 | 3 | 3 | 11 | 11 | 11 | 19 | 19 | 19 | 27 | 27 | 27 | 35 | 35 | 35 | 43 | 43 | 43 | hfB1 Standard 3 | hfB1 Standard 3 | hfB1 Standard 3 | | | |
| F | 51 | 51 | 51 | 59 | 59 | 59 | 67 | 67 | 67 | 75 | 75 | 75 | 83 | 83 | 83 | 91 | 91 | 91 | | | | | | |
| H | 4 | 4 | 4 | 12 | 12 | 12 | 20 | 20 | 20 | 28 | 28 | 28 | 36 | 36 | 36 | 44 | 44 | 44 | hFB1 Standard 4 | hFB1 Standard 4 | hFB1 Standard 4 | | | |
| G | 52 | 52 | 52 | 60 | 60 | 60 | 68 | 68 | 68 | 76 | 76 | 76 | 84 | 84 | 84 | 92 | 92 | 92 | | | | | | |
| I | 5 | 5 | 5 | 13 | 13 | 13 | 21 | 21 | 21 | 29 | 29 | 29 | 37 | 37 | 37 | 45 | 45 | 45 | hFB1 Standard 5 | hFB1 Standard 5 | hFB1 Standard 5 | | | |
| J | 53 | 53 | 53 | 61 | 61 | 61 | 69 | 69 | 69 | 77 | 77 | 77 | 85 | 85 | 85 | 93 | 93 | 93 | | | | | | |
| K | 6 | 6 | 6 | 14 | 14 | 14 | 22 | 22 | 22 | 30 | 30 | 30 | 38 | 38 | 38 | 46 | 46 | 46 | hFB1 Standard 6 | hFB1 Standard 6 | hFB1 Standard 6 | | | |
| L | 54 | 54 | 54 | 62 | 62 | 62 | 70 | 70 | 70 | 78 | 78 | 78 | 86 | 86 | 86 | 94 | 94 | 94 | | | | | | |
| M | 7 | 7 | 7 | 15 | 15 | 15 | 23 | 23 | 23 | 31 | 31 | 31 | 39 | 39 | 39 | 47 | 47 | 47 | hFB1 Standard 7 | hFB1 Standard 7 | hFB1 Standard 7 | | | |
| N | 55 | 55 | 55 | 63 | 63 | 63 | 71 | 71 | 71 | 79 | 79 | 79 | 87 | 87 | 87 | 95 | 95 | 95 | | | | | | |
| O | 8 | 8 | 8 | 16 | 16 | 16 | 24 | 24 | 24 | 32 | 32 | 32 | 40 | 40 | 40 | 48 | 48 | 48 | hFB1 Standard 8 | hFB1 Standard 8 | hFB1 Standard 8 | | | |
| P | 56 | 56 | 56 | 64 | 64 | 64 | 72 | 72 | 72 | 80 | 80 | 80 | 88 | 88 | 88 | 96 | 96 | 96 | | | | | | |

- Add 8µL of each DNA sample to designated wells for a total reaction volume of 20 µL.
- Seal the plate well with MicroAmp optical adhesive film (ThermoFisher # 4311971).
- To ensure reactants are at the bottom of the wells, briefly centrifuge the plate at 1,000g x 10 seconds.

#### *Running the qPCR assay*

- Load the sealed plate into the qPCR machine.

- Set cycling conditions according to the protocol your mastermix requires.

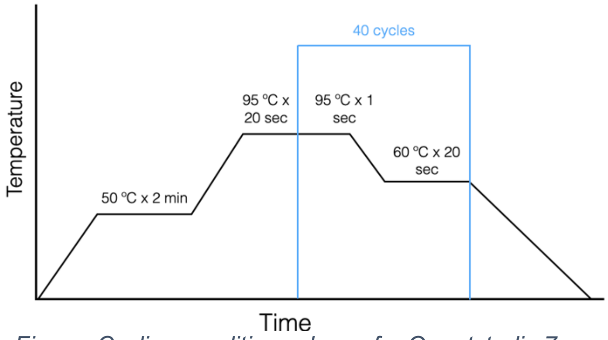


*Figure: Cycling conditions shown for Quantstudio 7 using TaqMan Universal Mastermix Fast (Life Technologies #4444965)*

- If using the TaqMan Universal MasterMix Fast (Life Technologies #4444965), cycling conditions are 50ºC for 2 minutes, followed by 95ºC for 20 seconds, then 40 cycles of 95ºC for 1 second followed by 60ºC for 20 seconds.
    - Total run time is about 40 minutes.

#### *qPCR data analysis and error checking*

- Compute the mean, standard deviation, and coefficient of variation (CV) for the cycle thresholds (Ct) across each triplicate for each sample.
- If the CV across the 3 triplicates is >2%, check the triplicates to see if any of the wells has a Ct that is >1 unit different from the other two, which could indicate a qPCR failure in that well.
    - In the case where amplification has failed in a well, remove the outlier.
- The average Ct for each sample should be used compute copies/µL.

### Appendix A: qPCR Reagents and MasterMix Recipe

#### qPCR Reagents

**Recipe for each well of qPCR**

| | |
|---|---|
| TaqMan Universal MasterMix Fast | 10 µL |
| ND1 Primers F+R | + 0.6 µL |
| ND1 Probe | + 0.4 µL |
| B2M Primers F+R | + 0.6 µL |
| B2M Probe | + 0.4 µL |
| **Total Reagent Volume** | **= 12 µL** |
| + Sample Volume | + 8 µL |
| **Total Reaction Volume** | **= 20 µL** |

The reagents used may depend on the qPCR equipment available, the genomic locations of interest, or other experimental parameters. Below, we provide a summary of possible reagents as well as other validated primers and probes.

**2x TaqMan Universal MasterMix Fast** (Life Tech #4444965)

**MicroAmp™ Optical 384-Well Reaction Plate with Barcode** (ThermoFisher # 4309849)

**MicroAmp™ Optical Adhesive Film** (ThermoFisher # 4311971)

**Validated mtDNA/nDNA Primer & Probe sequences** (idtDNA.com)

| Reference | qPCR Target | Sequences (5′→3′) |
|---|---|---|
| Trumpff C, et al.[3] | *mt-ND1*-F | GAGCGATGGTGAGAGCTAAGGT |
| | *mt-ND1*-R | CCCTAAAACCCGCCACATCT |
| | *mt-ND1*-Probe | HEX-CCATCACCCTCTACATCACCGCCC-3IABkFQ |
| | *B2m*-F | TCTCTCTCCATTCTTCAGTAAGTCAACT |
| | *B2m*-R | CCAGCAGAGAATGGAAAGTCAA |
| | *B2m*-Probe | FAM-ATGTGTCTGGGTTTCATCCATCCGACA-3IABkFQ |
| Ware, S, et al.[4] | *mt-ND1*-F | GAGCGATGGTGAGAGCTAAGGT |
| | *mt-ND1*-R | CCCTAAAACCCGCCACATCT |
| | *mt-ND1*-Probe | /5HEX/CCATCACCC/ZEN/TCTACATCACCGCCC-/3IABkFQ/ |
| | *B2m*-F | TCTCTCTCCATTCTTCAGTAAGTCAACT |
| | *B2m*-R | CCAGCAGAGAATGGAAAGTCAA |
| | *B2m*-Probe | /56-FAM/ATGTGTCTG/ZEN/GGTTTCATCCATCCGACA/3IABkFQ/ |

[3] Psychoneuroendocrinology. 2019 Aug;106:268-276. doi: 10.1016/j.psyneuen.2019.03.026. PMID: 31029929

[4] J Biol Chem. 2020 Nov 13;295(46):15677-15691. doi: 10.1074/jbc.RA120.015237. Epub 2020 Sep 8. PMID: 32900851; PMCID: PMC7667980.

| | | |
|---|---|---|
| Fazzini F, et al. [5] | *mt-tRNA$^{leu}$*-F | CACCCAAGAACAGGGTTTGT |
| | *mt-tRNA$^{leu}$*-R | TGGCCATGGGTATGTTGTTA |
| | *mt-tRNA$^{leu}$*-Probe | FAM-5′-TTACCGGGCTCTGCCATCT-BHQ1 |
| | *B2m*-F | TGCTGTCTCCATGTTTGATGTATCT |
| | *B2m*-R | TCTCTGCTCCCCACCTCTAAGT |
| | *B2m*-Probe | Yakima Yellow-5′-CAGGTTGCTCCACAGGTAGCTCTAG-BHQ1 |
| Rosa HS, et al.[6] | *hMito_F3* | CACTTTCCACACAGACATCA |
| | *hMito_R3* | TGGTTAGGCTGGTGTTAGGG |
| | *hB2M_F1* | TGTTCCTGCTGGGTAGCTCT |
| | *hB2M_R1* | CCTCCATGATGCTGCTTACA |

- Reconstitute primers in appropriate volume of nuclease free water to achieve 100µM stock concentration.
- Combine 120µL each of 100µM forward and reverse primers with 960 µL nuclease free water to achieve 10 µM working concentration.
- Store primers at -30 ºC or -80 ºC until use.
- Reconstitute probe in appropriate volume of nuclease free water to achieve 100µM stock concentration.
- Dilute 100 µM stock probe 20x to achieve 5µM working concentration.
- Store probes at -30$^{o}$C or -80$^{o}$C until use (avoid freeze-thaw) and protect probes from light.

[5] Sci Rep. 2018 Oct 18;8(1):15347. doi: 10.1038/s41598-018-33684-5. PMID: 30337569
[6] FASEB J. 2020 Jul 29. doi: 10.1096/fj.202000959RR. Epub ahead of print. PMID: 32729179.
